# A mouse model of Adams-Oliver Syndrome reveals defective Notch1 signaling in endothelial cells as a driver of pathogenesis

**DOI:** 10.1101/2025.03.20.644338

**Authors:** Alyssa F. Solano, Kristina Preusse, Brittany Cain, Rebecca Hotz, Parthav Gavini, Zhenyu Yuan, Benjamin Bowen, Gabrielle Maco, Hope Neal, Ellen K. Gagliani, Laura Southgate, Rhett A. Kovall, Raphael Kopan, Brian Gebelein

**Author notes:** Correspondence: Brian Gebelein, 3333 Burnet Avenue, MLC 7007, Cincinnati, OH 45229, Raphael Kopan, Ph.D., 3333 Burnet Avenue, MLC 7007, Cincinnati, OH 45229.

## Abstract

Adams-Oliver Syndrome (AOS) is a rare congenital disorder characterized by scalp, limb, and cardiovascular defects. While variants in the NOTCH1 receptor, DLL4 ligand, and RBPJ transcription factor have been implicated in AOS, the driving tissue types and molecular mechanisms by which these variants cause pathogenesis are unknown. Here, we used quantitative binding assays to show that AOS-associated RBPJ missense variants compromise DNA binding but not cofactor binding. These findings suggest that AOS-associated RBPJ variants do not function as loss-of-function alleles but instead act as dominant-negative proteins that sequester cofactors from DNA. Consistent with this idea, mice carrying an AOS-associated *Rbpj* allele develop dominant phenotypes that include lethality and cardiovascular defects in a *Notch1* heterozygous background, whereas *Notch1* and *Rbpj* compound heterozygous null alleles are well-tolerated. To facilitate studies into the key tissues driving AOS pathogenesis, we employed conditional genetics to isolate the contribution of the vasculature to the development of AOS-like phenotypes. Importantly, our studies show that expression of the *Rbpj* AOS allele in endothelial cells is both necessary and sufficient to cause lethality and cardiovascular defects. These data establish that defective Notch1 signaling in the vasculature is a key driver of pathogenesis in this AOS mouse model.

## Introduction

Adams-Oliver Syndrome (AOS) is a rare congenital condition primarily characterized by aplasia cutis congenita, which is a thinning and/or absence of the skin and skull tissue at the top of the head, and transverse terminal limb truncations that impact the hands and/or feet (1,2). AOS patients can also present with a wide range of cardiovascular defects that include congenital heart defects such as atrial and ventricular septal defects, valve anomalies, aortic and pulmonic stenosis, coarctation of the aorta, patent ductus arteriosus, persistent truncus arteriosus, pulmonary atresia, tetralogy of Fallot, and total or partial anomalous pulmonary venous drainage (3). Within the vasculature, a subset of AOS patients have defects such as cutis marmorata telangiectatica congenita (a congenital capillary malformation), portal vein agenesis, portal hypertension, esophageal varices, intracranial hemorrhages, and thrombosis (2). A smaller number of AOS patients have neurological defects such as microcephaly, ventricular dilation, corpus callosum hypoplasia, periventricular lesions, hypoplasia of optic nerves, visual deficits, cerebral palsy, epilepsy, spasticity, and cognitive impairment (2). Approximately 10% have intrauterine growth restriction (2). Hence, AOS patients experience a complex mixture of clinical phenotypes requiring a multidisciplinary approach to clinical management.

Molecular genetic studies have revealed that approximately 40% of AOS patients inherit variant alleles in one of six genes: *NOTCH1*, *DLL4*, *RBPJ*, *EOGT*, *DOCK6*, and *ARHGAP31* (2). AOS cases caused by variants in *NOTCH1*, *DLL4*, *RBPJ*, and *ARHGAP31* show an autosomal dominant inheritance pattern (4–7), while *EOGT* and *DOCK6* variants are inherited in an autosomal recessive manner (8,9). Of these six genes, four encode components of the Notch signaling pathway, including the receptor NOTCH1, the ligand DLL4, the transcription factor RBPJ, and the EGF domain-specific O-linked N-acetylglucosamine transferase EOGT, which post-translationally modifies Notch proteins (10). The remaining two genes encode proteins that regulate small GTPases, with *DOCK6* encoding a guanine nucleotide exchange factor and *ARHGAP31* encoding a Rho GTPase-activating protein (4,8). The relationship between the Notch pathway and the small GTPase regulators in AOS pathogenesis is currently unclear. However, patients with Notch pathway variants have a higher prevalence of cardiovascular defects than AOS patients without Notch pathway defects (49% vs. 13%), whereas AOS patients with pathogenic *DOCK6* variants have a higher prevalence of brain anomalies than other AOS patients (91% vs. 19%) (2). Overall, AOS pathogenesis remains poorly understood, and no disease-modifying therapies are available.

At the molecular level, the Notch signaling pathway converts extracellular ligand/receptor interactions between adjacent cells into functional changes in gene expression. Signaling is initiated when a ligand (DLL1, DLL3, DLL4, JAG1, or JAG2 in mammals) on a signal-sending cell binds to a transmembrane receptor (NOTCH1, NOTCH2, NOTCH3, or NOTCH4 in mammals) on the surface of the signal-receiving cell (10). Force generated during endocytosis of the ligand induces a conformation change in the receptor that allows proteolytic cleavage within the transmembrane region of NOTCH to release the Notch Intracellular Domain (NICD) from the cell surface (11). NICD then translocates to the nucleus and forms a transcriptional activation complex with the transcription factor RBPJ and the co-activator MAML to activate transcription of Notch target genes (10,12). However, RBPJ can also antagonize Notch signaling by recruiting corepressors to limit Notch target gene transcription (13–15).

Notch signaling is highly conserved across metazoans, and animal studies have revealed the Notch pathway regulates fundamental cellular processes that include cell fate specification, cell proliferation, and cell death (16,17). Throughout development, Notch signaling is iteratively used to regulate the morphogenesis of many organ systems, including the heart (18), vasculature (19), hematopoietic system (20), nervous system (21), and somite-derived organs (22). This is consistent with clinical studies showing that aberrant Notch signaling has been implicated in a wide array of health disorders that include AOS, bicuspid aortic valve disease, hypoplastic left heart syndrome, Alagille Syndrome, cerebral autosomal dominant arteriopathy with subcortical infarcts and leukoencephalopahty (CADASIL), Hajdu-Cheney Syndrome, spondylocostal dysostosis, and several types of cancer (23,24). How specific defects in the Notch pathway cause this broad array of disease is an active area of research.

Both due to the implication of Notch pathway genes in AOS and the observation of vascular changes in AOS patients (2,25), some have speculated that impaired vascular development during embryogenesis may drive the pathogenesis of the constellation of features observed in AOS (6,25–27). However, a vascular etiology for AOS has yet to be established, and the heart, skin/scalp, and limb defects found in AOS could alternatively be caused by defective Notch signaling in multiple cell types, including those in the skin, bone, and heart tissues (24). Unfortunately, the loss of a *Notch1* allele in mice is not sufficient to recapitulate AOS-like phenotypes, whereas the loss of a *Dll4* allele is so severe that heterozygotes rarely survive to birth due to catastrophic vascular defects (28,29). Tissue-specific conditional knockout of *Dll4* has been used to bypass this early lethality and model the impact of AOS-associated *Dll4* loss-of-function variants in mice (30), but the requirement for tissue-specificity limits the applications of this model. Thus, we currently lack a good mouse model of AOS to study pathogenesis.

Molecular genetic studies of AOS patients have uncovered frameshift and early truncation defects in *NOTCH1* and *DLL4* that are likely to render each allele null (2). These findings are consistent with dominant *NOTCH1* and *DLL4* variants creating loss-of-function alleles and *NOTCH1* and *DLL4* haploinsufficiency causing AOS (31). In contrast, all reported AOS-associated *RBPJ* variants are single-amino acid missense substitutions; no frameshift or nonsense *RBPJ* variants have been identified that would encode obvious null alleles. To better understand the molecular mechanism by which AOS-associated *RBPJ* variants impact Notch signaling, our group previously leveraged an existing *Drosophila melanogaster* line with an E137V missense mutation in *Suppressor of Hairless* (*Su(H)*, fly ortholog of *RBPJ*) that is analogous to an AOS-associated variant in human *RBPJ* at residue E63 (32). Intriguingly, a single copy of the *Su(H)*^E137V^ allele was sufficient to induce a wing nicking phenotype, which is not seen in flies with a single copy of a *Su(H)* null allele. Moreover, the *Su(H)*^E137V^ allele was also sufficient to dramatically enhance the loss of sensory bristle phenotype associated with haploinsufficiency of the *Hairless (H)* corepressor, whereas a single copy of a *Su(H)* null allele suppresses this phenotype (32,33). We subsequently used biochemical methods to show that the Su(H)^E137V^ protein selectively decreased DNA binding, while binding to NICD and the Hairless corepressor was unaffected. Taken together, these findings support the model that the *Su(H)^E137V^* allele encodes a dominant-negative protein that dysregulates Notch signaling by sequestering the NICD activator and Hairless corepressor proteins from DNA regulatory elements.

In this study, we use quantitative DNA binding assays to show that all six AOS-associated *RBPJ* alleles encode proteins with defective DNA binding activity, but with differing degrees of severity, ranging from a 3-fold decrease relative to wild-type Rbpj to a complete loss in DNA binding. To assess how such alleles would impact mammalian development, we used genome editing to make two different mouse models that encode AOS-associated Rbpj variants with ∼3-fold (Rbpj^S358R^) and ∼6-fold (Rbpj^E89G^) decreased DNA binding activity. Characterization of these mice reveal that, while each allele compromises the Notch pathway, these alleles are not sufficient to generate dominant phenotypes in an otherwise wild-type mouse line. However, mice that are compound heterozygous for a *Notch1* null allele and the *Rbpj^E89G^* allele had dramatically decreased viability and showed pronounced vascular and heart defects. In contrast, compound heterozygous mice with both *Notch1* and *Rbpj* null alleles were born at normal Mendelian ratios and showed no gross morphological defects. These findings are consistent with the AOS-associated *Rbpj* variants encoding dominant-negative proteins and not null alleles. Since an *Rbpj* null allele is well-tolerated in mice, we designed a novel conditional genetic approach to demonstrate that expression of the *Rbpj^E89G^* dominant-negative allele in endothelial cells is both necessary and sufficient to induce lethality due to vascular and heart-related defects. These studies provide novel insights into how defective Notch signaling in the vasculature is a key driver of AOS pathogenesis in mice and thereby serves as a useful model to study human AOS pathogenesis.

## Results

### AOS-associated Rbpj variants reduce DNA but not cofactor binding

RBPJ has a highly conserved core consisting of four distinct regions including an N-terminal domain, beta-trefoil domain, interdomain linker, and C-terminal domain (Figure 1A). In the human ortholog (NM_005349.4), residues 57-67 and 165-170 in the N-terminal domain and 192-197 in the beta-trefoil domain directly interact with DNA (Figure 1B and Kovall and Hendrickson, 2004). To date, six likely deleterious RBPJ variants have been reported in AOS, all of which are single-amino acid missense substitutions that alter highly conserved residues (Y60C, E63G, R65G, F66V, K169E, and S332R; Figure 1A) (2,5). Five of these missense variants occur within the RBPJ DNA binding domain, with S332R falling within a conserved short interdomain linker region (Figure 1A). Consistent with the locations of these point mutations, prior studies characterized the DNA binding properties of a subset of these RBPJ disease variants (E63G and K169E) and found decreased DNA binding (5). These studies led to the prediction that the AOS-associated RBPJ variants cause disease by behaving as loss-of-function alleles due to decreased DNA binding.

**Figure 1:**
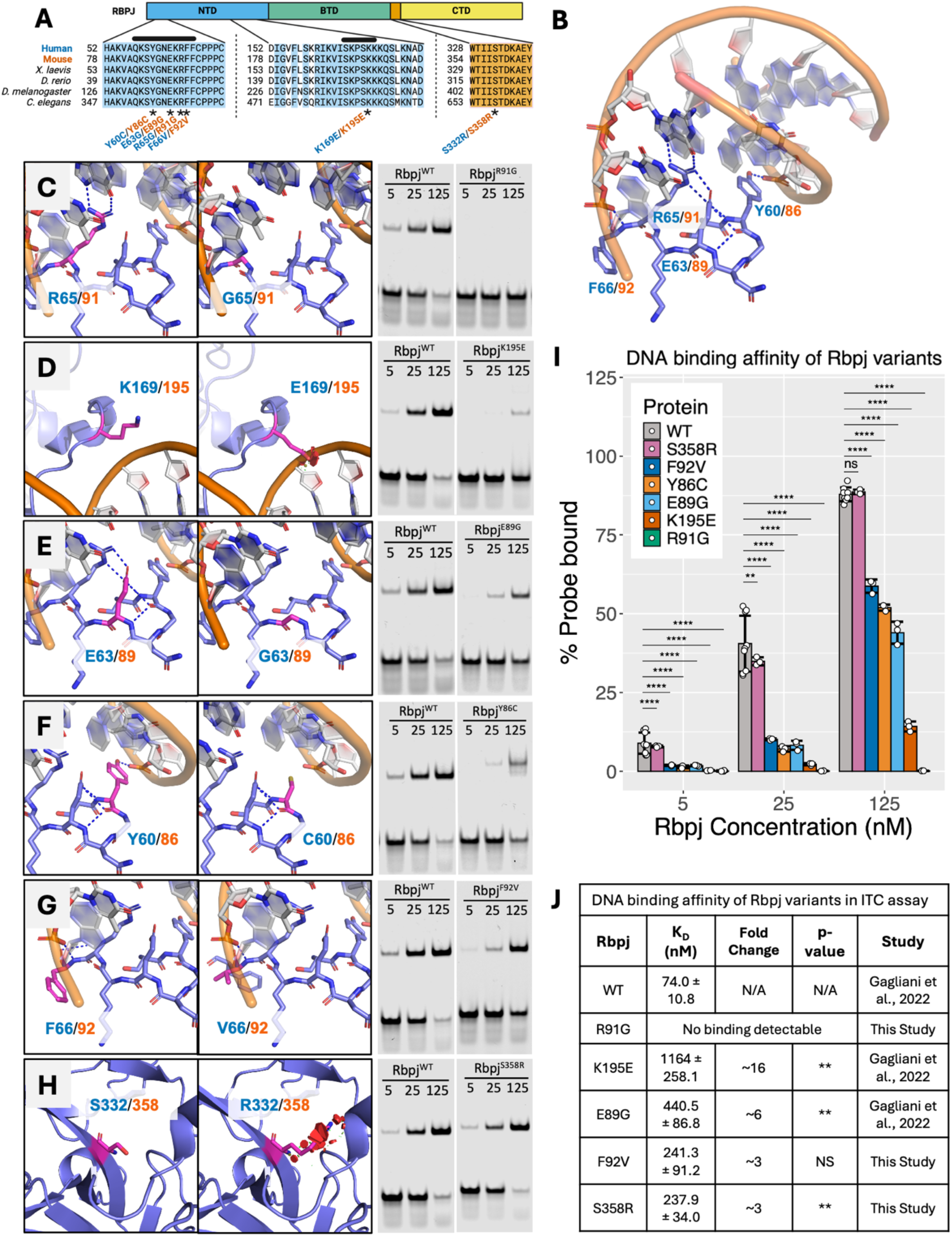
AOS-associated RBPJ variants impair DNA binding affinity. A) Domain map and sequence alignment of RBPJ orthologs. Conserved residues are highlighted, and AOS-associated variants (*) are denoted by human (blue) and mouse (orange) residue numbers. Black bars indicate DNA-binding regions. NTD = N-terminal domain. BTD = beta-trefoil domain. CTD = C-terminal domain. Created with *BioRender*. B) Structure of wild-type (WT) Rbpj on DNA with residues mutated in AOS denoted by human (blue) and mouse (orange) residue numbers. C-H) PyMol models of structural changes and representative comparative EMSAs of AOS-associated RBPJ variants. Dashed lines within each model denote DNA-residue or residue-residue polar interactions and red discs indicate steric clash. EMSAs were performed using equimolar concentrations (5, 25, and 125 nM) of wild-type mouse Rbpj and the R91G (C), K195E (D), E89G (E), Y86C (F), F92V (G), and S358R (H) variants on a DNA probe with a single high-affinity Rbpj binding site. I) Graph showing the quantification of probe depletion for each variant across triplicate EMSAs (see Figure S1). A one-way ANOVA with Tukey post-hoc correction was used to compare wild-type Rbpj to each variant. J) Tabulated ITC data measuring DNA binding affinity of Rbpj variants. Fold change is calculated relative to wild-type Rbpj. A two-tailed t-test was used to compare the dissociation constant (K_D_) of wild-type Rbpj to each variant. [*] p < 0.05. [**] p < 0.01. [***] p < 0.001. [****] p < 0.0001. N/A = not applicable. NS = not significant.

To determine if all RBPJ AOS variants negatively impact DNA binding and directly compare the DNA binding activities of each variant, we performed electrophoretic mobility shift assays (EMSAs) and isothermal titration calorimetry (ITC) assays using DNA probes with a high affinity Rbpj binding site and purified AOS-associated Rbpj variants within the context of the mouse Rbpj protein (Figure 1 and Supplemental Figures S1 and S2). Moreover, we modeled each variant in the context of the known RBPJ/DNA structure to compare with our DNA binding results and better understand the molecular nature of each defect (Figure 1C-H). Note, we previously performed and reported ITC assays to assess the DNA binding affinity of wild-type Rbpj and the Rbpj^E89G^ and Rbpj^K195E^ AOS variants. We included that data here along with new EMSA data for comparative purposes and cited the original source of this data as appropriate. Collectively, these studies revealed two key findings: first, all variants significantly decreased DNA binding compared to wild-type Rbpj; and second, the variants’ impact on DNA binding fell across a spectrum of severity (Figure 1C-J and Supplemental Figures S1 and S2). Below, we describe the impact of each variant.

The most severe variant was Rbpj^R91G^, which completely abolished DNA binding in both EMSAs (Figure 1C and 1I) and ITC assays (Figure 1J and Supplemental Figure 2A). This finding was conserved across all Rbpj concentrations tested and is congruent with the R91G residue change predicted to abolish direct polar interactions with both DNA and the adjacent amino acid E89 (Figure 1C). Almost as severe was the Rbpj^K195E^ variant, which significantly compromised DNA binding in EMSAs (Figure 1D and 1I) and decreased binding ∼16-fold compared to wild-type Rbpj in ITC assays (Figure 1J and Supplemental Figure 2A). Consistent with the dramatic loss in DNA binding, the K195E change introduces electrostatic repulsion and steric clashing within a region involved in direct binding to the DNA backbone (Figure 1D).

The Rbpj^E89G^ and Rbpj^Y86C^ variants decreased DNA binding to a similar extent in EMSAs (Figure 1E, 1F, and 1I). ITC assays further showed that the Rbpj^E89G^ substitution resulted in an approximate ∼6-fold loss in DNA binding activity relative to wild-type Rbpj (Figure 1J and Supplemental Figure 2A). Consistent with these findings, the E89G residue change is predicted to abolish polar interactions with adjacent residues including both Y86 and R91. Unfortunately, we were unable to purify sufficient Rbpj^Y86C^ protein to perform ITC assays. Moreover, the Rbpj^Y86C^/DNA complex migrated slower than those of wild-type Rbpj and all other tested variants in EMSAs, even though these proteins were similar in size in SDS gels (Supplemental Figure S1B). Since the Rbpj^Y86C^ variant introduces a new Cys residue, we treated the protein samples with different reducing agents and performed EMSAs but did not observe a change in this slower migration pattern (Supplemental Figure S1C). While it is currently unclear why the Y86C substitution results in a slower migrating band, the similar loss of affinity observed by the Rbpj^Y86C^ and Rbpj^E89G^ proteins in EMSAs is consistent with the structural analysis showing that Y86C is predicted to disrupt both polar and nonpolar interactions with DNA (Figure 1F).

In comparison, the last two variants, Rbpj^F92V^ and Rbpj^S358R^, resulted in weaker but still significant decreases in DNA binding in EMSAs compared to wild-type Rbpj (Figure 1G and 1H). ITC assays confirmed a ∼3-fold decrease in DNA binding affinity for each of these variants relative to wild-type Rbpj (Figure 1J and Supplemental Figure 2A). The modest impact on DNA binding is consistent with the S358R variant residing in a linker region that does not directly contact DNA. However, this variant is predicted to induce steric clashing with surrounding residues (Figure 1H), and thereby could cause protein folding changes that result in decreased DNA binding. The F92V variant is not predicted to change existing polar interactions or introduce steric clashing. However, F92 appears to make substantial nonpolar interactions with the DNA backbone that the smaller V92 residue may not fully recapitulate (Figure 1G), which could explain its modest reduction in DNA binding activity. Taken together, these DNA binding assays show that all Rbpj AOS variants negatively impact DNA binding but do so to varying degrees.

The above DNA binding findings support the idea that AOS-associated *Rbpj* alleles encode defective transcription factors that fail to properly bind DNA. In addition to binding DNA, RBPJ directly recruits the NICD signaling molecule to activate gene expression and co-repressors to inhibit gene expression. We previously showed that two of the AOS variants, Rbpj^E89G^ and Rbpj^K195E^, do not significantly alter their affinity for the NICD co-activator or the SHARP co-repressor protein (32). We similarly found that the Rbpj^F92V^ variant binds both NICD and SHARP with similar affinities as wild-type Rbpj and that the Rbpj^R91G^ variant binds to NICD with a similar affinity as wild-type Rbpj in ITC assays (Supplemental Figure 2B, 2C, and Table S1). Since the Y86C variant is found within the same region of the DNA binding domain as E89G, R91G, and F92V, which is far from the NICD and SHARP interaction regions, this variant is also unlikely to alter binding to these cofactor proteins. However, because S358R is located within a linker region not directly associated with DNA or cofactor binding, we also tested Rbpj^S358R^ in quantitative ITC assays with purified NICD and SHARP proteins (Supplemental Figure 2B and 2C). Importantly, these data show that Rbpj^S358R^ binds to NICD and SHARP with the same affinity as the wild-type Rbpj protein (Table S1). Thus, these data support the model that all known RBPJ variants associated with AOS negatively impact DNA binding, but not cofactor binding, consistent with these Rbpj variants acting as dominant-negative proteins that sequester cofactors away from wild-type Rbpj and off DNA.

**Figure 2:**
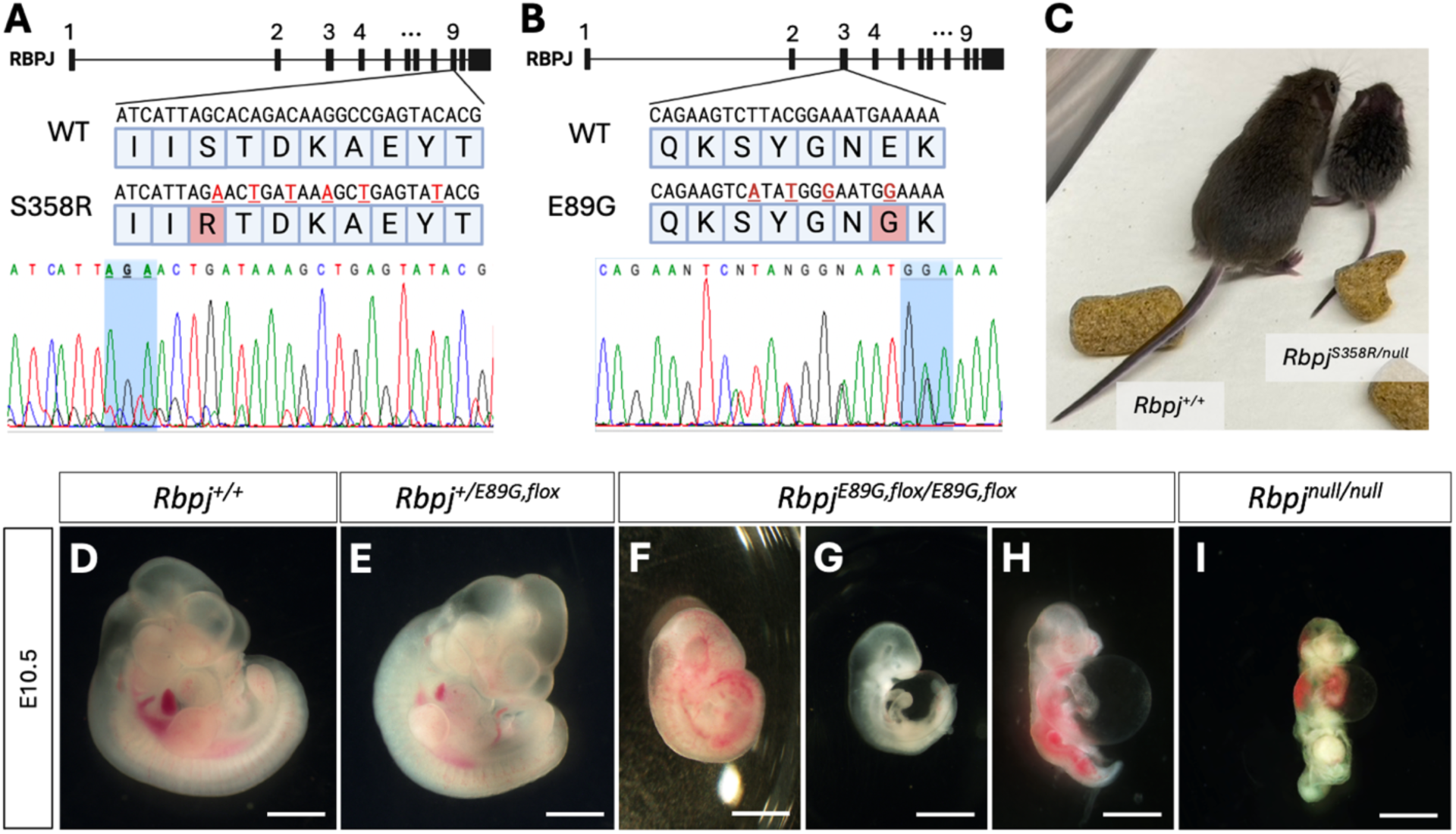
Generation of AOS-associated Rbpj variant mouse models reveals impaired animal growth and development. A) (*top*) Schematic of mouse *Rbpj*, detailing the region of exon 9 that encodes S358 and the donor sequence used to introduce an S358R substitution. Created with *BioRender*. (*bottom*) Confirmation of mouse genotype by Sanger sequencing. The codon for S358/R358 is highlighted. B) (*top*) Schematic of mouse *Rbpj*, detailing the region of exon 3 that encodes E89 and the donor sequence used to introduce an E89G substitution. Created with *BioRender*. (*bottom*) Confirmation of mouse genotype by Sanger sequencing. The codon for E89/G89 is highlighted. C) Image showing that a typical P17 *Rbpj^S358R/null^* hemizygote (right) is much smaller than its *Rbpj^+/+^* littermate (left). D-I) Stereoscope images of mice at embryonic day 10.5 (E10.5) show that *Rbpj^E89G,flox/E89G,flox^* homozygotes (F-H) display growth retardation, hemorrhage, pallor, and/or pericardial edema of variable severity. *Rbpj^null/null^* homozygotes (I) show similar defects, albeit more severe. Scale bar = 1 mm.

### *Rbpj^E89G^* and *Rbpj^S358R^* mouse models reveal that phenotypic severity correlates with loss in DNA binding affinity

To make mouse models carrying AOS-associated *Rbpj* alleles, we used CRISPR/Cas9 gene editing to engineer two analogous *Rbpj* mutations that have been associated with AOS. We chose to model the *Rbpj^S358R^* and *Rbpj^E89G^* variants based on their mild (∼3-fold loss) and moderate (∼6-fold loss) impacts on DNA binding affinity, respectively, to avoid potential heterozygote lethality in a mouse carrying a severe variant. To introduce the S358R substitution (analogous to human S332R), we used a donor sequence to replace part of exon 9 of mouse *Rbpj* (Figure 2A). We similarly introduced the E89G substitution (analogous to human E63G) using a donor sequence to replace part of exon 3 (Figure 2B). In both cases, silent mutations were also included in the donor sequences to introduce restriction enzyme sites that facilitate genotyping (see methods for details), and each variant was subsequently confirmed by sequencing (Figure 2A-B). Note, while the *Rbpj^S358R^* variant was generated on a wild-type *Rbpj* allele, we created the *Rbpj^E89G^* variant on the well-characterized floxed *Rbpj^flox^* allele (35). Our rationale for making the *Rbpj^E89G^* variant on the floxed allele is that Cre recombinase can be used to convert the dominant-negative *Rbpj^E89G,flox^* allele into an *Rbpj^null^* allele in select tissues of heterozygous mice that still have a non-floxed wild-type *Rbpj* allele (i.e. *Rbpj^+/E89G,flox^*).

To determine the impact of these two *Rbpj* alleles on mouse viability, we first assessed offspring for deviation from expected Mendelian ratios using chi-squared tests. From these studies, we found that *Rbpj^+/S358R^* heterozygous and *Rbpj^S358R/S358R^* homozygous mice were viable and occurred at expected ratios (Table 1). Moreover, these mice did not show any gross morphological defects, although the *Rbpj^S358R/S358R^* mice were initially smaller than littermates but were of normal size by postnatal week 5 (Supplemental Figure 3). These data indicate that the *Rbpj^S358R^* allele is well-tolerated in mice. To further test this idea, we crossed *Rbpj^S358R/S358R^* mice with mice carrying an *Rbpj* null allele (*Rbpj^+/null^*) and found that *Rbpj^S358R/null^* hemizygotes had significantly reduced viability (Table 1) and surviving offspring were much smaller than their littermates (Figure 2C). These studies are consistent with the Rbpj^S358R^ variant having a modest impact on DNA binding (∼3-fold loss compared to wild-type Rbpj). Thus, the *Rbpj^S358R^* allele behaves as a weak hypomorph in mice.

**Table 1:** Impact of *Rbpj* variants on pre- and post-natal mouse viability.

| Cross | | Number of Pups | Genotype of Interest | Expected (%) | Observed (%) | $\chi^2$ p-value |
| --- | --- | --- | --- | --- | --- | --- |
| Rbpj<sup>+/S358R</sup> x Rbpj<sup>+/S358R</sup> |  | 135 | Rbpj<sup>S358R/S358R</sup> | 33.75 (25%) | 26 (19%) | NS |
| Rbpj<sup>+/S358R</sup> x Rbpj<sup>+/null</sup> |  | 67 | Rbpj<sup>S358R/null</sup> | 16.75 (25%) | 6 (9.0%) | 0.0079 |
| Rbpj<sup>+/E89G, flox</sup> x Rbpj<sup>+/E89G, flox</sup> |  | 166 | Rbpj<sup>E89G, flox/E89G, flox</sup> | 41.5 (25%) | 0 (0.0%) | 5.40 x 10<sup>-13</sup> |
|  | Rbpj<sup>+/E89G, flox</sup> x Rbpj<sup>+/E89G, flox</sup> (E10.5) | 81 | Rbpj<sup>E89G, flox/E89G, flox</sup> | 20.25 (25%) | 14 (17.3%) | 0.0171 |
| Rbpj<sup>+/E89G, flox</sup> x Rbpj<sup>+/S358R</sup> |  | 75 | Rbpj<sup>E89G, flox/S358R</sup> | 18.75 (25%) | 1 (1.3%) | 1.71 x 10<sup>-5</sup> |

Except where noted, mice were genotyped postnatally. For each noted genotype of
interest, the expected and observed numbers and percentages are reported with Chi
squared analysis ( $\chi^2$ ) used to assess for significant deviations from Mendelian ratios. NS
= not significant.

We similarly assessed the impact of the *Rbpj^E89G,flox^* allele on viability and found that while heterozygous mice (*Rbpj^+/E89G,flox^*) were viable and lack gross morphological defects, no *Rbpj^E89G,flox/E89G,flox^* homozygous mice were observed among live offspring (Table 1). To assess when during embryogenesis *Rbpj^E89G,flox/E89G,flox^* homozygotes perish, we performed timed collections at embryonic day 10.5 (E10.5). While *Rbpj^+/E89G,flox^* embryos resemble their wild-type littermates (Figure 2D and 2E), we observed a lower than expected frequency of *Rbpj^E89G,flox/E89G,flox^* embryos (Table 1) and all homozygous embryos were much smaller than their littermates (Figure 2F-H). Moreover, these embryos showed a range of morphological defects consistent with decreased Notch signaling that included widespread areas of hemorrhage (Figure 2F, n = 4/8), pericardial edema (Figure 2G and 2H, n = 6/8), pallor (Figure 2G, n = 3/8), and lack of complete axial rotation (Figure 2H, n = 3/8). The pericardial edema and lack of complete axial rotation are reminiscent of *Rbpj^null/null^* embryos (Figure 2I), although the *Rbpj^E89G,flox/E89G,flox^* embryos fare slightly better than *Rbpj^null/null^* embryos and the *Rbpj^E89G,flox/E89G,flox^* embryos express normal levels of Rbpj protein relative to wild-type Rbpj (Supplemental Figure 4). Lastly, we crossed *Rbpj^+/E89G,flox^* mice with *Rbpj^+/S358R^* mice and observed a dramatic loss of viability in offspring with both the *Rbpj^S358R^* and *Rbpj^E89G,flox^* alleles (*Rbpj^S358R/E89G,flox^*, Table 1). Taken together, these data show that that the Rbpj^E89G^ variant, which has an ∼6-fold decrease in DNA binding activity, causes more severe phenotypes in mice than the Rbpj^S358R^ variant with ∼3-fold loss in DNA binding.

### A compound heterozygous mouse model carrying *Rbpj^E89G^* and *N1^null^* AOS alleles shows vascular and heart phenotypes

Our mouse data with the *Rbpj^S358R^* and *Rbpj^E89G,flox^* alleles, which are analogous to known AOS-associated *RBPJ* alleles, reveals that neither of these alleles is sufficient to cause AOS-like phenotypes as heterozygotes in mice. In contrast, patients heterozygous for these alleles develop dominant AOS phenotypes, although the *RBPJ^S332R^* allele was found in only a single patient and has incomplete penetrance (2). These findings are consistent with prior studies showing differences in sensitivity to Notch pathway alleles between mice and humans. For example, *NOTCH1* haploinsufficiency can cause human disease such as AOS and aortic valve disease (23), whereas a *Notch1* (*N1*) null allele is well-tolerated in a heterozygous state in mice (31,36). Interestingly, however, a family with AOS was found to have compound heterozygous mutations in both *RBPJ* and *NOTCH1* alleles (2). Hence, we crossed *Rbpj^+/E89G,flox^* mice with mice heterozygous for either a *N1* null allele that deletes amino acids 1056-2049, which encode several EGF repeats, the transmembrane region, and the Ankyrin repeats (*N1^tm1Con^* (36), referred to here as *N1^null^*), or a *N1* null allele that deletes the promoter and exon 1 (*N1^tm2Agt^*(37), referred to here as *N1^gKO^*; see Methods for more details of each allele). Importantly, we observed a dramatic loss of viability in both *N1^+/null^;Rbpj^+/E89G,flox^* and *N1^+/gKO^;Rbpj^+/E89G,flox^* compound heterozygous mice (Table 2), and the surviving postnatal mice generally failed to thrive. Intriguingly, we found that a subset of the *N1^+/gKO^;Rbpj^+/E89G,flox^* compound heterozygotes, which had considerable C57/BL6 in their background, had obvious gross morphological skin/scalp defects (Figure 3A-B). These findings raise the possibility of genetic background contributing to the skin/scalp defect. Hence, in this study, we focus on identifying the mechanisms of embryonic lethality, which was observed with both *N1* alleles and on an outbred background.

**Figure 3:**
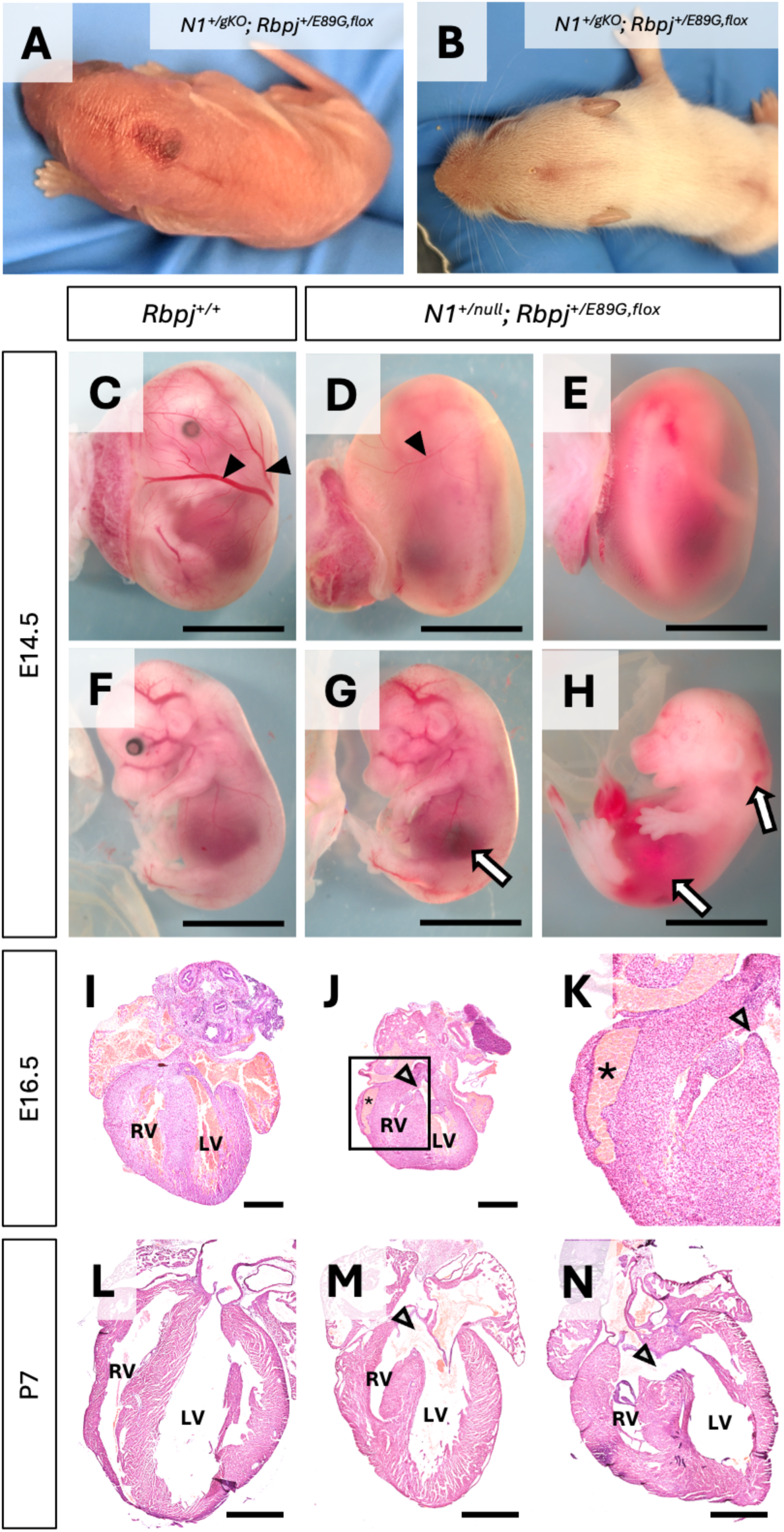
*N1^+/null^; Rbpj^+/E89G,flox^* embryos display cardiovascular phenotypes. A-B) Representative images of dorsal midline skin lesions in P0 (A) and P11 (B) *N1^+/gKO^; Rbpj^+/E89G,flox^* mice. C-E) Representative image of E14.5 embryos within their yolk sac for wild-type (*Rbpj^+/+^*) and *N1^+/null^; Rbpj^+/E89G,flox^* genotypes. Note, the compound heterozygous embryos have reduced or absent yolk sac vasculature (filled arrowheads). F-H) Representative images of E14.5 embryos for wild-type (*Rbpj^+/+^*) and *N1^+/null^; Rbpj^+/E89G,flox^* genotypes. Note, areas of hemorrhage (arrows) are observed in E14.5 *N1^+/null^; Rbpj^+/E89G,flox^* embryos but not in control embryos. I-K) Representative images of E16.5 H&E stained heart sections from wild-type (*Rbpj^+/+^*) and *N1^+/null^; Rbpj^+/E89G,flox^* genotypes. The left ventricles (LV) and right ventricles (RV) are labeled and arrowheads highlight ventricular septal defects in *N1^+/null^; Rbpj^+/E89G,flox^* heart, whereas asterisks highlight dilated coronary vessels. The box in (J) outlines the region shown at higher magnification in (K). L-N) Representative images of P7 H&E stained heart sections from wild-type (*Rbpj^+/+^*) and *N1^+/null^; Rbpj^+/E89G,flox^* genotypes. The left ventricles (LV) and right ventricles (RV) are labeled and arrowheads highlight ventricular septal defects in *N1^+/null^; Rbpj^+/E89G,flox^* hearts. Scale bars are 0.5 cm (C-H), 0.5 mm (I-J), and 1 mm (L-N).

**Table 2:** Impact of *Rbpj* variants on pre- and post-natal mouse viability in Notch1-sensitized backgrounds.

| Cross | Number of Pups | Genotype of Interest | Expected (%) | Observed (%) | $\chi^2$ p-value |
| --- | --- | --- | --- | --- | --- |
| $Rbpj^{+/E89G, flox} \times N1^{+/gKO}$ | 89 | $Rbpj^{+/E89G, flox}; N1^{+/gKO}$ | 22.25 (25%) | 3 (3.4%) | $2.92 \times 10^{-7}$ |
| $Rbpj^{+/E89G, flox} \times N1^{+/null}$ | 255 | $Rbpj^{+/E89G, flox}; N1^{+/null}$ | 63.75 (25%) | 10 (3.9%) | $3.82 \times 10^{-13}$ |
| $Rbpj^{+/E89G, flox} \times N1^{+/null}$ (E16.5) | 214 | $Rbpj^{+/E89G, flox}; N1^{+/null}$ | 53.5 (25%) | 21 (9.8%) | $4.27 \times 10^{-6}$ |
| $Rbpj^{+/E89G, flox} \times N1^{+/null}$ (E14.5) | 48 | $Rbpj^{+/E89G, flox}; N1^{+/null}$ | 12 (25%) | 8 (16.7%) | NS |
| $Rbpj^{+/E89G, flox} \times N1^{+/null}$ (E10.5) | 22 | $Rbpj^{+/E89G, flox}; N1^{+/null}$ | 5.5 (25%) | 5 (22.7%) | NS |
| $Rbpj^{+/null} \times N1^{+/gKO}$ | 29 | $Rbpj^{+/null}; N1^{+/gKO}$ | 7.25 (25%) | 8 (27.6%) | NS |
| $Rbpj^{+/null} \times N1^{+/null}$ | 91 | $Rbpj^{+/null}; N1^{+/null}$ | 22.75 (25%) | 19 (20.9%) | NS |
| $Rbpj^{+/E89G, flox} \times N2^{+/lacZ}$ | 106 | $Rbpj^{+/E89G, flox}; N2^{+/lacZ}$ | 26.5 (25%) | 18 (17.0%) | NS |

Except where noted, mice were genotyped postnatally. For each noted genotype of interest, the expected and observed numbers and percentages are reported with Chi squared analysis ( $\chi^2$ ) used to assess for significant deviations from Mendelian ratios. NS = not significant.

We next tested the specificity of the genetic interactions between the *N1* null alleles and the *Rbpj^E89G^* AOS variant allele that cause lethality by performing two additional genetic tests. First, we crossed each *N1* null allele with mice carrying an *Rbpj* null allele and found that neither *N1^+/null^;Rbpj^+/null^* nor *N1^+/gKO^;Rbpj^+/null^* were significantly underrepresented (Table 2). Moreover, unlike the *N1* and *Rbpj^+/E89G,flox^* compound heterozygotes that showed morphological defects and failed to thrive, the *N1^+/null^;Rbpj^+/null^* and *N1^+/gKO^;Rbpj^+/null^* compound heterozygous mice were indistinguishable in appearance from littermate controls. These data show that the decreased viability observed in the *N1* and *Rbpj^+/E89G,flox^* compound heterozygous mice was due to the presence of the *Rbpj^E89G,flox^* allele and not simply due to loss of a wild-type *Rbpj* allele. Second, we crossed mice carrying the *Rbpj^+/E89G,flox^* allele into a *Notch2 (N2)*-sensitized background and observed expected numbers of *N2^+/lacZ^;Rbpj^+/E89G,flox^* compound heterozygotes that showed no gross morphological defects (Table 2). Thus, the *Rbpj^E89G^* allele genetically interacts with *N1* null alleles to cause decreased viability but not with a *N2* null allele. These mouse data are consistent with clinical findings showing that RBPJ variants have been shown to cause a N1-like syndrome (AOS) but not a N2-like syndrome (Alagille) (38).

The decreased viability and failure of *N1^+/null^;Rbpj^+/E89G,flox^* mice to thrive made it difficult to obtain sufficient animals to perform a comprehensive quantitative analysis of tissues using postnatal mice. To better define the cause of increased lethality in *N1^+/null^;Rbpj^+/E89G,flox^* compound heterozygous embryos, we next genotyped embryos from timed harvests at E10.5, E14.5, and E16.5 to assess the time of embryonic demise. These experiments revealed there is a gradual decrease in *N1^+/null^;Rbpj^+/E89G,flox^* compound heterozygous embryos that becomes significant by E16.5 (Table 2). Moreover, gross morphological analysis of these embryos revealed several vascular phenotypes that included reductions in the yolk sac vasculature (Figure 3C-E) and hemorrhages (Figure 3F-H). Since heart defects are commonly found in both humans and mice with Notch pathway defects, we also analyzed E16.5 hearts and observed malformations that included ventricular septal defects (VSDs) and dilated coronary vessels in *N1^+/null^;Rbpj^+/E89G,flox^* compound heterozygous embryos (Figure 3I-K, note we quantify these data through additional genetic crosses below). Consistent with these data, analysis of the hearts from the relatively few postnatal day 7 (P7) *N1^+/null^;Rbpj^+/E89G,flox^* mice revealed that one third also had VSDs (2 of 6, Figure 3J-L). While *N1* variants in humans have been associated with bicuspid valve disease, we did not observe obvious valve abnormalities in the hearts of either E16.5 nor P7 *N1^+/null^;Rbpj^+/E89G,flox^* animals. Taken together, these data show that *N1^+/null^;Rbpj^+/E89G,flox^* compound heterozygous mice show increased embryonic lethality that is potentially caused by cardiovascular defects.

### Conditional removal of the AOS-associated *Rbpj^E89G,flox^* allele from only the vasculature rescues cardiovascular phenotypes

Two key pieces of evidence have led to the hypothesis that AOS is largely a vascular disease. First, AOS patients with *N1*, *DLL4*, and *RBPJ* variants often suffer cardiovascular defects (2). Second, mouse and zebrafish studies have shown that N1 and Dll4 signaling are critical regulators of vascular development (39,40). To test this hypothesis, we developed an AOS “rescue” mouse model that takes advantage of conditional genetics by using the *Tie2-Cre^Ywa^* line to specifically recombine floxed alleles in the developing endothelial cells (ECs), which form the inner lining of the vasculature, endocardial cells, and hematopoietic stem cells (HSCs). By crossing *N1^+/null^*;*Tie2-Cre^+/Ywa^* mice with *Rbpj^+/E89G,flox^* mice, Cre-mediated recombination converts the floxed *Rbpj^E89G,flox^* allele into an *Rbpj^null^* allele in heterozygous ECs and HSCs that still encode a wild-type *Rbpj^+^* allele (see schematics in Figure 4A). Importantly, since *N1^+/null^;Rbpj^+/null^* mice occur in expected numbers (Table 2) and do not show overt phenotypes, this mouse model explicitly tests if expressing the dominant-negative *Rbpj^+/E89G,flox^* allele within the vascular and blood systems is required (i.e. necessary) to induce morbidity in a *N1^+/null^* background (Figure 4A). Consistent with this idea, we found that *N1^+/null^;Rbpj^+/E89G,flox^* compound heterozygous mice with *Tie2-cre* had significantly enhanced viability compared to *N1^+/null^;Rbpj^+/E89G,flox^* littermates that lack *Tie2-cre* (Table 3). Moreover, the postnatal *N1^+/null^;Rbpj^+/E89G,flox^* mice with *Tie2-cre* were indistinguishable from control littermates, whereas the *N1^+/null^;Rbpj^+/E89G,flox^* mice without *Tie2-cre* generally failed to thrive (Supplementary Figure 5). Thus, *Tie2-cre* can significantly rescue the embryonic lethality seen in *N1^+/null^;Rbpj^+/E89G,flox^* compound heterozygous mice by converting the *Rbpj^+/E89G,flox^* AOS allele into a *Rbpj^+/null^* allele within the developing vasculature.

**Figure 4:**
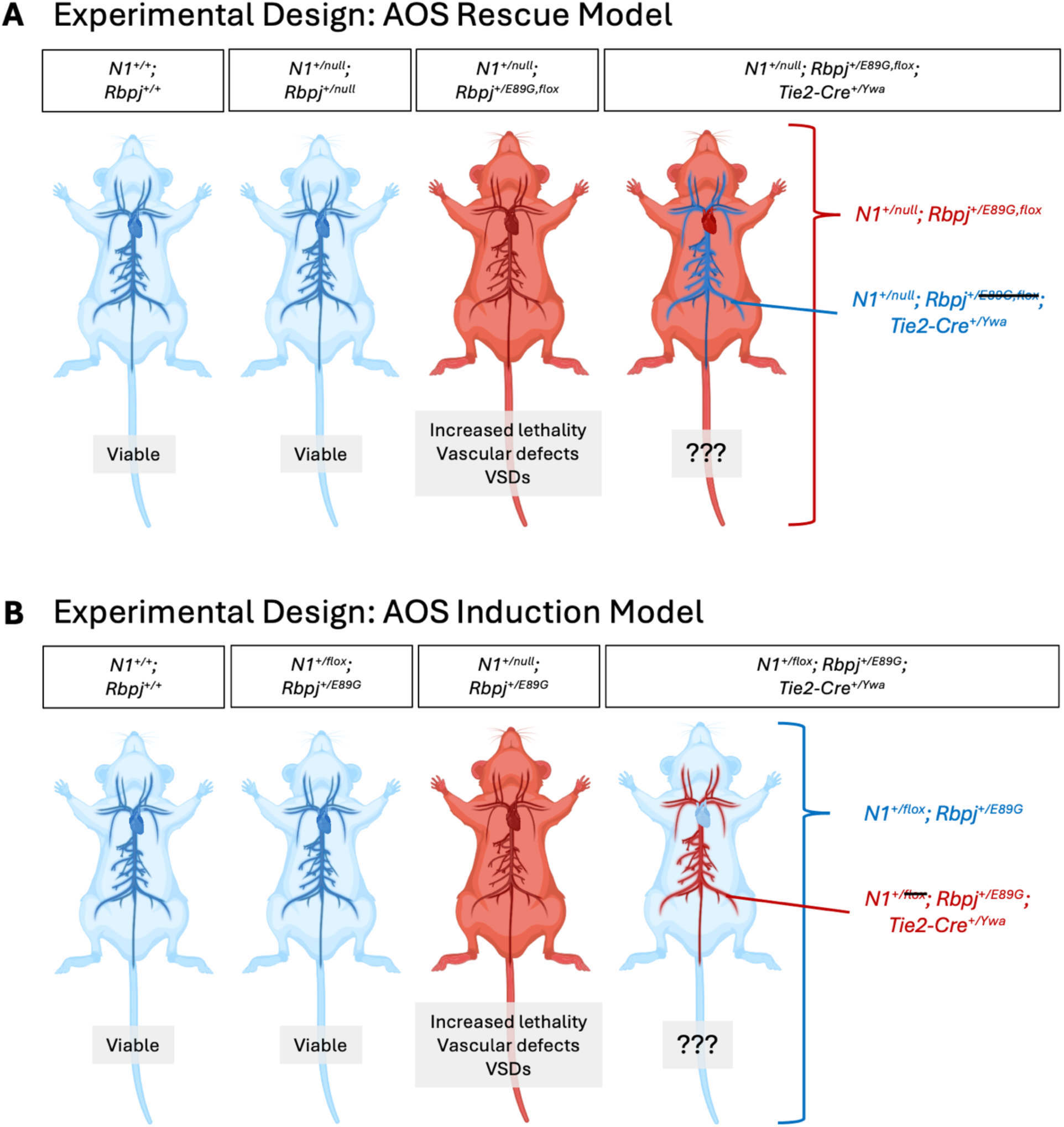
Selective removal or induction of a *N1^+/null^; Rbpj^+/E89G^* genotype in the vasculature using Tie2-cre-mediated recombination tests the contribution of the vasculature to the development of AOS-like cardiovascular phenotypes. A) Schematics of AOS rescue model. Both wild-type (*N1^+/+^; Rbpj^+/+^*) and *N1^+/null^; Rbpj^+/null^* mice are viable and without overt defects. Mice that globally express the *N1^+/null^; Rbpj^+/E89G,flox^* genotype have reduced viability, vascular defects, and heart defects (see Table 3 and Figure 3). A proposed mouse that recombines *N1^+/null^; Rbpj^+/E89G,flox^* to *N1^+/null^; Rbpj^+/null^* only in the vasculature via expression of Cre recombinase under the vascular-specific promoter *Tie2* would test the necessity of the mutation’s presence in the vasculature for the development of AOS-like phenotypes. B) Schematics of AOS induction model. Both wild-type (*N1^+/+^; Rbpj^+/+^*) and *N1^+/flox^; Rbpj^+/E89G^* mice are viable and without overt defects (see Table 4). Based on data from *N1^+/null^; Rbpj^+/E89G,flox^* mice (see Table 3 and Figure 3), mice that globally express the *N1^+/null^; Rbpj^+/E89G^* genotype are predicted to have reduced viability, vascular defects, and heart defects. A proposed mouse that recombines *N1^+/flox^; Rbpj^+/E89G^* to *N1^+/cKO^; Rbpj^+/E89G^* only in the vasculature via expression of Cre recombinase under the vascular-specific promoter *Tie2* would test the sufficiency of the mutation’s presence in the vasculature for the development of AOS-like phenotypes. Created with *BioRender*.

**Table 3:** Impact of *Rbpj* variants in the vasculature on pre- and post-natal mouse viability in Notch-sensitized backgrounds.

| Cross | Number of Pups | Genotype of Interest | Expected (%) | Observed (%) | $\chi^2$ p-value |
| --- | --- | --- | --- | --- | --- |
| *Rbpj*<sup>+/E89G,flox</sup> x *N1*<sup>+/-null</sup>, *Tie2-Cre*<sup>+/-Ywa</sup> | 128 | *Rbpj*<sup>+/E89G,flox</sup>; *N1*<sup>+/-null</sup> | 16<br>(12.5%) | 5<br>(3.9%) | 0.0232 |
| *Rbpj*<sup>+/E89G,flox</sup> x *N1*<sup>+/-null</sup>, *Tie2-Cre*<sup>+/-Ywa</sup> | 128 | *Rbpj*<sup>+/E89G,flox</sup>; *N1*<sup>+/-null</sup>; *Tie2-Cre*<sup>+/-Ywa</sup> | 16<br>(12.5%) | 11<br>(8.6%) | NS |
| *Rbpj*<sup>+/E89G,flox</sup> x *N1*<sup>+/-null</sup>, *Tie2-Cre*<sup>+/-Ywa</sup> (E16.5) | 104 | *Rbpj*<sup>+/E89G,flox</sup>; *N1*<sup>+/-null</sup> | 13<br>(12.5%) | 3<br>(2.9%) | 0.0139 |
| *Rbpj*<sup>+/E89G,flox</sup> x *N1*<sup>+/-null</sup>, *Tie2-Cre*<sup>+/-Ywa</sup> (E16.5) | 104 | *Rbpj*<sup>+/E89G,flox</sup>; *N1*<sup>+/-null</sup>; *Tie2-Cre*<sup>+/-Ywa</sup> | 13<br>(12.5%) | 12<br>(11.5%) | NS |

Except where noted, mice were genotyped postnatally. For each noted genotype of interest, the expected and observed numbers and percentages are reported with Chi squared analysis ( $\chi^2$ ) used to assess for significant deviations from Mendelian ratios. NS = not significant.

**Table 4:** Impact of *Rbpj* variants in the vasculature on pre- and post-natal mouse viability in Notch-sensitized backgrounds.

| Cross | Number of Pups | Genotype of Interest | Expected (%) | Observed (%) | $\chi^2$ p-value |
| --- | --- | --- | --- | --- | --- |
| *Rbpj*<sup>+/E89G</sup>; Tie2-Cre<sup>+/Ywa</sup><br>x *N1*<sup>flox/flox</sup> | 66 | *Rbpj*<sup>+/E89G</sup>; *N1*<sup>+/flox</sup> | 16.5<br>(25%) | 18<br>(27.3%) | NS |
| *Rbpj*<sup>+/E89G</sup>; Tie2-Cre<sup>+/Ywa</sup><br>x *N1*<sup>flox/flox</sup> | 66 | *Rbpj*<sup>+/E89G</sup>; *N1*<sup>+/flox</sup>;<br>Tie2-Cre<sup>+/Ywa</sup> | 16.5<br>(25%) | 1<br>(1.52%) | 3.87 x 10<sup>-05</sup> |
| *Rbpj*<sup>+/E89G</sup>; Tie2-Cre<sup>+/Ywa</sup> x *N1*<sup>flox/flox</sup><br>(E16.5) | 116 | *Rbpj*<sup>+/E89G</sup>; *N1*<sup>+/flox</sup> | 29<br>(25%) | 44<br>(37.9%) | NS |
| *Rbpj*<sup>+/E89G</sup>; Tie2-Cre<sup>+/Ywa</sup> x *N1*<sup>flox/flox</sup><br>(E16.5) | 116 | *Rbpj*<sup>+/E89G</sup>; *N1*<sup>+/flox</sup>;<br>Tie2-Cre<sup>+/Ywa</sup> | 29<br>(25%) | 13<br>(11.2%) | 0.0208 |

Except where noted, mice were genotyped postnatally. For each noted genotype of interest, the expected and observed numbers and percentages are reported with Chi squared analysis ( $\chi^2$ ) used to assess for significant deviations from Mendelian ratios. NS = not significant.

Because we obtain relatively few *N1^+/null^;Rbpj^+/E89G,flox^* postnatal mice without *Tie2-cre*, we quantified the impact of conditionally converting the *Rbpj^+/E89G,flox^* allele into an *Rbpj^+/null^* allele in embryos. To do so, we first performed timed embryonic collections at E14.5 and E16.5 and assessed for embryonic viability and yolk sac vascular defects. Consistent with our postnatal analysis, we found that *Tie2-cre* was sufficient to rescue lethality of *N1^+/null^;Rbpj^+/E89G,flox^* embryos at E16.5, whereas *N1^+/null^;Rbpj^+/E89G,flox^* embryos without *Tie2-cre* were significantly underrepresented at this timepoint (Table 3). Moreover, visual analysis of the embryonic yolk sac at both E14.5 and E16.5 revealed that *Tie2-cre* could significantly rescue the observed vascular defects of *N1^+/null^;Rbpj^+/E89G,flox^* embryos (Figure 5A-J). For example, while 7 of 9 E14.5 *N1^+/null^;Rbpj^+/E89G,flox^* embryos had reduced or absent yolk sac vasculature, 0 of 6 E14.5 *N1^+/null^;Rbpj^+/E89G,flox^;Tie2-cre^+/Ywa^* embryos and none of the other control littermates showed diminished yolk sac vasculature (Figure 5A-E and 5P). A similar rescue in yolk sac vasculature was observed in *Tie2-cre* positive *N1^+/null^;Rbpj^+/E89G,flox^* embryos at E16.5 (Figure 5F-J and 5P). Thus, these data show that conditionally converting the *Rbpj^+/E89G,flox^* allele into an *Rbpj^+/null^* allele with *Tie2-cre* was sufficient to rescue both viability and the yolk sac vasculature defects in *N1^+/null^* heterozygous embryos. Intriguingly, comparative analysis between embryonic timepoints revealed that the observed penetrance of yolk sac vasculature defects in the absence of *Tie2-cre* was significantly decreased at E16.5 (∼33%) compared to E14.5 (∼78%) in *N1^+/null^;Rbpj^+/E89G,flox^* embryos (p = 0.046). This decreased penetrance of yolk sac vascular defects in older embryos correlates well with the embryonic viability data showing a decrease in the proportion of *N1^+/null^;Rbpj^+/E89G,flox^* embryos from E14.5 to E16.5 (Table 2). Hence, these data suggest that those E14.5 embryos with severe yolk sac phenotypes are likely to perish prior to E16.5 and that conditionally deleting the *Rbpj^+/E89G,flox^* allele using *Tie2-cre* can rescue this phenotype and embryonic lethality.

**Figure 5:**
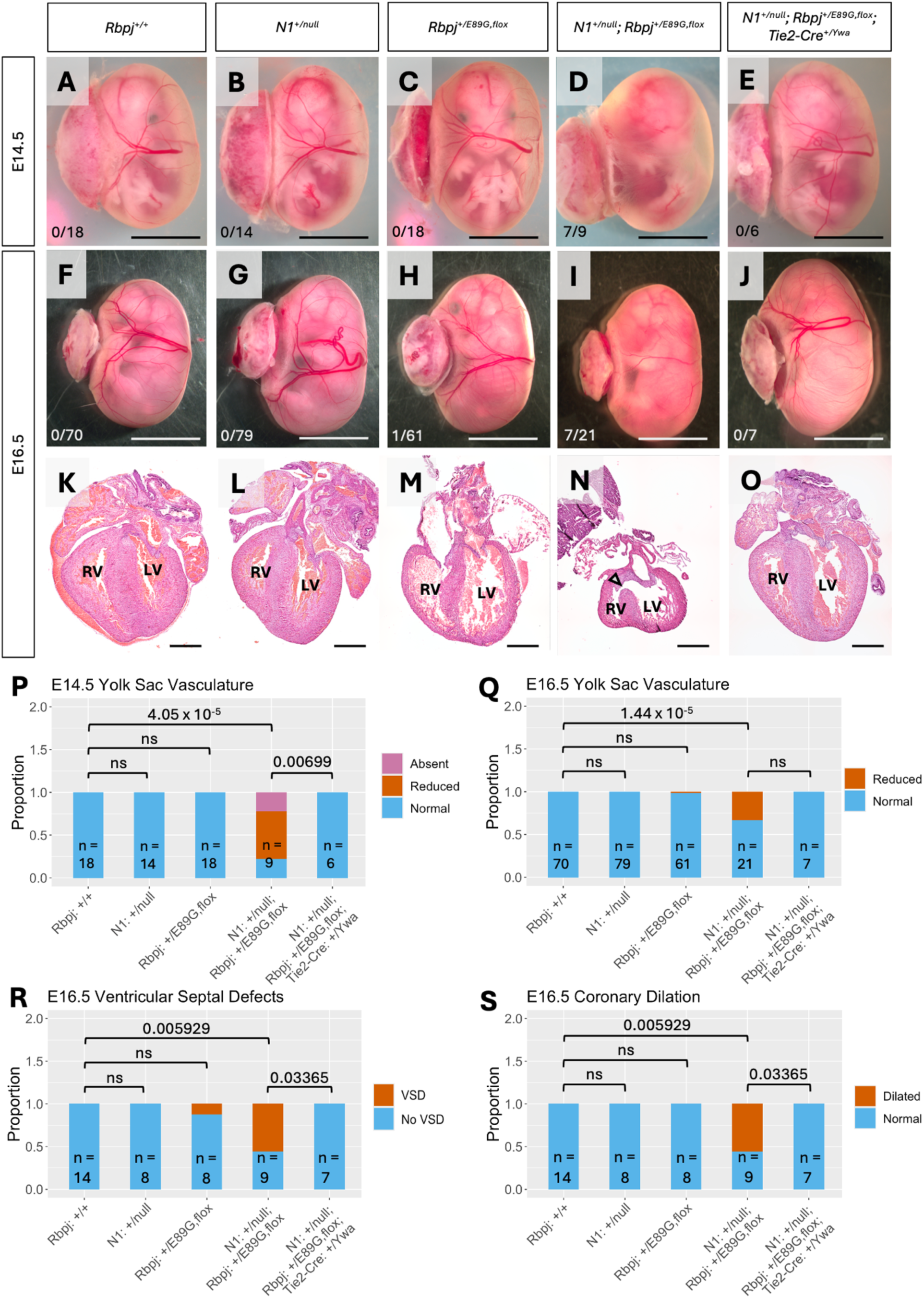
Conditional removal of *Rbpj^E89G^* from the vascular endothelium rescues vascular and heart phenotypes in mice. A-J) Representative image of E14.5 embryos (A-E) and E16.5 embryos (F-J) within their yolk sac for wild-type (*Rbpj^+/+^*), *N1*^+/*null*^, *Rbpj*^+/*E89G,flox*^, *N1*^+/*null*^;*Rbpj*^+/*E89G,flox*^, and *N1*^+/*null*^;*Rbpj*^+/*E89G,flox*^;*Tie2-cre^Ywa^* genotypes. Note, only the *N1^+/null^;Rbpj^+/E89G,flox^* embryos have reduced or absent yolk sac vasculature. The ratio of affected individuals to total individuals is listed in the lower left corner of each panel. K-O) Representative images of E16.5 H&E stained heart sections. The left ventricles (LV) and right ventricles (RV) are labeled and an arrowhead highlights a ventricular septal defect in the *N1^+/null^; Rbpj^+/E89G,flox^* heart. P-S) Visualization of the proportion of embryos with P-Q) yolk sac vasculature defects, R) ventricular septal defects, and S) dilated coronary vessels. p-values calculated with Fisher’s exact test are noted on the graphs. ns = not significant.

To further assess for possible vascular defects, we harvested and immunostained the skin vasculature from the forelimb and scalp regions of E14.5 embryos using a CD31 antibody to label ECs. Analysis of the forelimb tissues for both percent vascularized area and branch point density did not reveal significant differences in vascular development across genotypes (Supplemental Figure 6A-D). In addition, we analyzed tip cell numbers within the scalp skin vasculature at E14.5, a timepoint at which sprouting angiogenesis is still actively occurring at the top of the skull and did not observe any obvious changes in tip cell numbers across genotypes (Supplemental Figure 6E-I). Thus, while significant defects in the yolk sac vasculature are observed in the *N1^+/null^;Rbpj^+/E89G,flox^* compound heterozygotes, we did not observe widespread microvascular defects within the embryonic skin.

Next, we assessed if *Tie2-cre* could similarly rescue the observed embryonic heart defects seen in *N1^+/null^;Rbpj^+/E89G,flox^* compound heterozygotes (see Figure 3). Unlike wild-type embryos (Figure 5K), *N1^+/null^* single heterozygotes (Figure 5L), and *Rbpj^+/E89G,flox^* single heterozygotes (Figure 5M), the *N1^+/null^;Rbpj^+/E89G,flox^* compound heterozygotes showed significant heart defects at E16.5 that included VSDs (5 of 9, Figure 5N and 5R) and coronary vessel dilation (5 of 9, Figure 5S). In contrast, we did not observe any of these phenotypes in the *N1^+/null^;Rbpj^+/E89G,flox^*;*Tie2-Cre^+/Ywa^* embryos, suggesting that the heart defects in the *N1^+/null^;Rbpj^+/E89G,flox^* embryos are due to compromised Notch1 signaling in the developing endothelium and endocardium cells. Altogether, these results show that the expression of the AOS-associated dominant-negative Rbpj protein in the vascular endothelium is necessary for the development of cardiovascular phenotypes.

### Selective induction of *N1^+/cKO^;Rbpj^+/E89G^* compound heterozygosity in the vasculature is sufficient to cause embryonic lethality and cardiovascular phenotypes

The AOS rescue mouse model described above reveals that expression of the *Rbpj^E89G^* allele in the developing vascular system is required (i.e. necessary) to induce significant morbidity in *N1^+/null^* heterozygous mice. To test if *N1^+/null^;Rbpj^+/E89G^* compound heterozygous alleles within the developing vasculature are sufficient to induce morbidity, we modified our conditional genetic approach to create an AOS induction model (see schematics in Figure 4B). For this mouse model, we first used genome editing to remake the *Rbpj^E89G^* AOS variant on a non-floxed *Rbpj* allele. We subsequently generated *Rbpj^+/E89G^;Tie2-cre^+/Ywa^* mice and crossed these to *N1^flox/flox^* mice (41) to generate *N1^+/flox^;Rbpj^+/E89G^* offspring with and without *Tie2-cre*. Cre recombinase conditionally removes the promoter and exon 1 of the *N1^flox^* allele; we refer to this recombined allele as *N1^cKO^*. Note, *Tie2-cre* in this model will selectively induce *N1^+/cKO^;Rbpj^+/E89G^* compound heterozygosity within endothelial cells (and in the endocardium and HSCs) of mice that otherwise have cells with two copies of *N1* (i.e. *N1^+/flox^;Rbpj^+/E89G^*) (Figure 4B). Excitingly, *Notch1^+/flox^;Rbpj^+/E89G^;Tie2-Cre^+/Ywa^* mice occur significantly less often than their littermates, suggesting prenatal demise (Table 4). Moreover, consistent with previously described results (Figure 3 and 5) E16.5 *N1^+/flox^;Rbpj^+/E89G^;Tie2-cre^+/Ywa^* embryos had both significantly reduced yolk sac vasculature (Figure 6A-D) and increased incidences of major hemorrhage (Figure 6E-H) compared to littermates. Additionally, VSDs were observed in *N1^+/flox^;Rbpj^+/E89G^;Tie2-cre^+/Ywa^* embryonic hearts, but not in those of control littermates (3 of 7, Figure 6I-K). Taken together, these results show that carrying *N1^+/null^;Rbpj^+/E89G^* compound heterozygosity in the vasculature is sufficient for the developmental of cardiovascular phenotypes.

**Figure 6:**
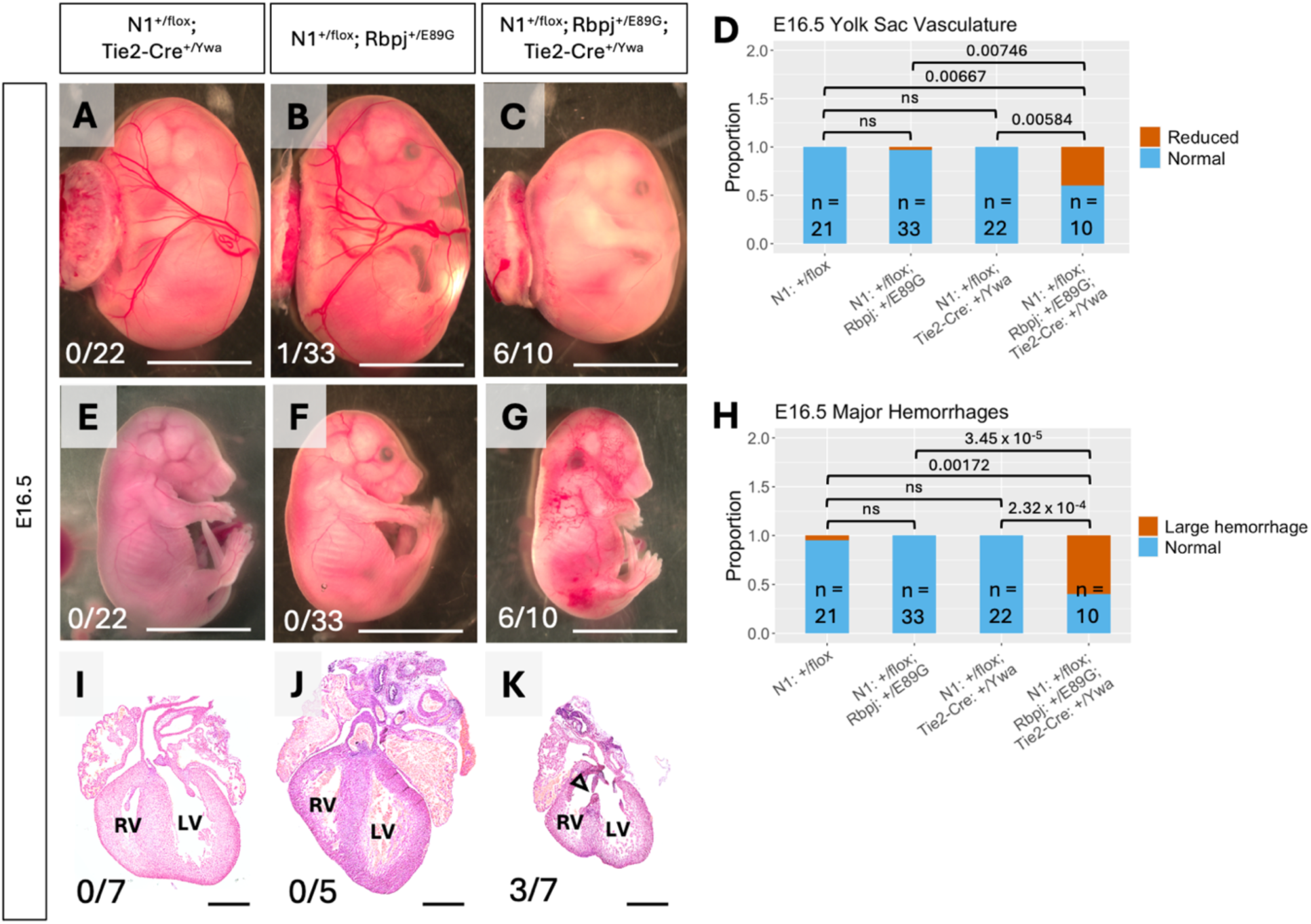
Conditional removal of one copy of *Notch1* from the vascular endothelium of *Rbpj^+/E89G^* mice induces vascular and heart phenotypes. E16.5 *N1^+/flox^; Rbpj^+/E89G^; Tie2-Cre^+/Ywa^* embryos have reduced yolk sac vasculature (A-D), increased frequency of hemorrhage (E-H), and VSDs (I-K). The left ventricles (LV) and right ventricles (RV) are labeled, and an arrowhead highlights a ventricular septal defects in the *N1^+/flox^; Rbpj^+/E89G^; Tie2-cre^Ywa^* heart. The ratio of affected individuals to total individuals is listed in the lower left corner of each panel. Scale bars are 0.5 cm (A-C and E-G) and 0.5 mm (I-K). ns = not significant.

## Discussion

In this study, we investigated the molecular and tissue-specific mechanisms underlying how AOS-associated RBPJ variants cause pathogenesis. At the molecular level, we used quantitative DNA binding and protein-protein interaction assays to show that all AOS-associated RBPJ variants tested to date have reduced binding to DNA but not to the transcriptional co-activator NICD or the transcriptional co-repressor SHARP. These findings suggest that AOS-associated RBPJ variants act as dominant-negative alleles in heterozygous cells by competing for cofactors with wild-type RBPJ and sequestering them off DNA. The idea that AOS-associated RBPJ variants do not act as loss-of-function null alleles is further supported by genetic studies in *Drosophila*, mice, and humans. We previously found that *Drosophila* with an analogous AOS mutation in the fly Rbpj homologue (*Su(H)*) develop dominant Notch phenotypes that are not observed in flies heterozygous for a *Su(H)* null allele (32). Here, we similarly found that mice heterozygous for the *Rbpj^E89G^* AOS allele suffer embryonic lethality and cardiovascular defects in a sensitized *N1* background, whereas mice compound heterozygous for *N1* and an *Rbpj* null allele occur in normal Mendelian ratios and suffer no obvious gross morphological defects. Lastly, molecular genetic studies of AOS patients to date have identified six missense variants that decrease DNA binding, whereas no nonsense or frameshift mutations have been identified that would render the *RBPJ* allele into a null allele (2,5). Moreover, a seventh AOS variant that also impacts the R65 residue (R65T) was recently reported on ClinVar (VCV001803755.1) and this variant is likely to negatively impact DNA binding in a manner similar to the R65G variant. Combined, these molecular and genetic data support the idea that AOS-associated *RBPJ* alleles encode dominant-negative proteins that dysregulate Notch signaling. Interestingly, however, even though *RBPJ* null alleles have not been implicated in AOS, they are severely underrepresented in the Genome Aggregation Database (pLI = 1; gnomAD v4.1.0) suggesting RBPJ haploinsufficiency is likely deleterious in humans (42). Thus, future studies are needed to determine what impact *RBPJ* haploinsufficiency has on human development.

Our comparative quantitative DNA binding studies between six AOS RBPJ variants revealed that, while each significantly compromised DNA binding, they do so to different degrees. These findings predict that the RBPJ variants that more strongly decrease DNA binding will result in greater dysregulation of the Notch pathway and worse outcomes. Consistent with this idea, mice with the Rbpj^E89G^ variant that decreases DNA binding 6-fold resulted in more severe phenotypes than mice with the Rbpj^S358R^ variant that decreases DNA binding 3-fold. Similarly, prior *Drosophila* studies found that the *Su(H)^T4^* allele that compromises DNA binding ∼5-fold resulted in more severe Notch pathway dysregulation and phenotypes compared to the *Su(H)^O5^* allele that encodes a protein with ∼3.5-fold decreased DNA binding (32). While the rarity of AOS in humans makes it impossible to perform a comprehensive comparison between variant DNA binding activity and clinical severity, it is interesting to note that the two variants with the weakest impact on DNA binding were found to either have incomplete penetrance (*RBPJ^S332R^*) or was only found in patients that carried both an *RBPJ^F66V^* allele and a rare missense *N1* allele (2). In contrast, the other RBPJ variants, which all impact DNA binding at least 6-fold, have not been associated with any other Notch pathway alleles, and to our knowledge all patients with a copy of these alleles have AOS phenotypes.

Through a combination of genome editing to make mice encoding AOS-associated RBPJ variants and conditional genetics to manipulate Notch pathway alleles, we generated a novel experimental model to identify the defective N1 signaling tissues that contribute to pathogenesis. Our approach takes advantage of the fact that only mice heterozygous for both a *N1* null allele and the AOS-associated *Rbpj^E89G^* allele (*N1^+/null^;Rbpj^+/E89G^*) suffer significant pathological phenotypes. Through tissue-specific cre-mediated recombination, we developed two conditional genetic approaches that either selectively remove the *Rbpj^E89G,flox^* allele in an otherwise *N1^+/null^* background or selectively induce *N1^+/null^;Rbpj^+/E89G^* compound heterozygous genotypes in the desired tissue (here, the developing vasculature; Figure 4). Since embryonic analysis of *N1^+/null^;Rbpj^+/E89G^* compound heterozygotes revealed significant lethality by E16.5 with embryos showing severe yolk sac vascular defects, hemorrhages, dilated coronary vessels and ventricular septal defects (VSDs), we selected the *Tie2-cre* line to manipulate either the *Rbpj^E89G,flox^* allele or a *N1^flox^* allele in each mouse model. Importantly, we found that *Tie2-cre*, which is expressed in endothelial and endocardial cells, can significantly rescue embryonic lethality and the observed vascular and heart defects by deleting the *Rbpj^E89G,flox^* allele in a *N1* heterozygous background. *Tie2-cre* can also significantly cause embryonic lethality and the observed vascular and heart defects by inducing *N1* heterozygosity in the presence of a *Rbpj^E89G^* allele. These data strongly support the idea that the *Rbpj^E89G^* allele causes pathological defects by dysregulating N1 signaling in endothelial cells. Moreover, these data demonstrate how novel conditional strategies can be applied to define the tissue-specific mechanisms of dominant-negative, disease-causing alleles with unclear etiology.

Our finding that the majority, if not all Rbpj^E89G^-mediated pathogenesis is due to compromised Notch signaling in endothelial cells is consistent with the vast literature showing that N1 and Dll4 signaling, two genes specifically associated with AOS, are critical regulators of vascular development. These findings lead to another question: What aspect of vascular development is compromised by the Rbpj^E89G^ variant? Notch signaling regulates many aspects of vascular development (43), including endothelial cell fate decisions (44), endothelial cell proliferation (45,46), arteriovenous specification (47,48), tip-stalk specification (49,50), mural cell patterning (51–55), and vascular integrity (52,54). The vascular defects observed in *N1^+/null^;Rbpj^+/E89G,flox^* mice include a paucity of yolk sac vessels and increased frequency of hemorrhage during late embryogenesis, suggesting both that vessels are not developing in sufficient quantity and that vascular integrity is compromised. Since prior studies showed that N1/Dll4 signaling is essential for sprouting angiogenesis by regulating the specification of appropriate numbers of tip versus stalk cell fates, we analyzed the vascular front in skin preparations from the E14.5 scalp but did not observe any obvious differences in tip cell numbers. In addition, we analyzed the embryonic forelimb skin vascular networks for changes in vascularized area and branch point density and found no difference between *N1^+/null^;Rbpj^+/E89G,flox^* and wild-type embryos. These studies indicate that the *N1^+/null^;Rbpj^+/E89G^* compound heterozygotes do not show a dramatic alteration in sprouting angiogenesis. However, additional studies with better temporal control using this novel mouse model and tamoxifen-inducible Cre lines are needed to provide a comprehensive assessment of how the *Rbpj^E89G^* allele impacts sprouting angiogenesis in an experimentally tractable tissue like the postnatal retina.

Notch signaling is also required for normal cardiac morphogenesis, as endocardial Notch activity contributes to development of the heart’s valves, chambers, and outflow tract (56). Consistent with these prior findings, *N1^+/null^;Rbpj^+/E89G,flox^* mice have abnormal cardiac morphology, most commonly membranous VSDs and dilated coronary vessels. These VSDs likely result from impaired growth or fusion of the endocardium with the cardiac neural crest-derived outflow tract septum (57). Dilated coronary vessels may be secondary to the heart failing (58). The lack of abnormal valve morphology in our mouse model is evidence that endocardial cells have successfully undergone epithelial-mesenchymal transition to form the endocardial cushions, a Notch-dependent process (59). This suggests that only some Notch-dependent processes are interrupted in this mouse model, perhaps due to variation in Notch dose sensitivity between different processes.

While this study focused on using conditional approaches with the *N1^+/null^;Rbpj^+/E89G,flox^* mouse model to define the mechanisms underlying embryonic lethality and cardiovascular defects, we were unable to assess if similar mechanisms underly skin/scalp and limb defects, two widely regarded hallmarks of AOS in human patients. In fact, in all our mouse studies, we never observed any obvious limb defects. However, we did observe skin/scalp defects with one of the *N1* alleles (*N1^gKO^*), which had considerable C57/BL6 in its genetic background. These findings raise the possibility that the N1-dependent skin/scalp lesions are sensitive to genetic background, which would be consistent with prior studies in mice showing that Notch-dependent phenotypes can vary based on genetic background (28). Thus, comparative studies are needed using inbred mice carrying conditional *N1* and *Rbpj^E89G^* alleles to isolate the role of genetic background and test whether the observed skin/scalp lesions are due defective N1 signaling in endothelial and/or other cell types using tissue-specific Cre lines.

Lastly, an unanswered question in the Notch field and human disease is how do variants in *RBPJ*, which is the sole transcription factor downstream of all NOTCH ligands and receptors, cause a N1/DLL4 associated syndrome (AOS) but not a N2/JAG1 associated syndrome (Alagille)? Consistent with these clinical data, we found that the *Rbpj^E89G^* allele in mice genetically interacts with *N1* haploinsufficiency to cause early embryonic lethality due to vascular and heart defects, whereas compound heterozygous mice for *Rbpj^E89G^* and a *N2* null allele are born at expected Mendelian ratios without any obvious morphological defects. While additional studies are needed to assess if the *Rbpj^E89G^* allele compromises N2-sensitive cell types to a sub-pathological degree, these data suggest that the *Rbpj^E89G^* allele preferentially compromises N1-over N2-dependent processes. Importantly, the mice generated in this study provide a new experimental model to test this hypothesis and explore potential mechanisms underlying how RBPJ AOS alleles broadly impact Notch biology.

## Methods

### Sex as a biological variable

AOS occurs in both males and females without obvious sex-bias (2,5,6). Nevertheless, we examined male and female mice and observed similar changes in viability in both sexes (see Supporting Data Values file for the sex of mice included in postnatal viability assays). Hence, we did not consider sex as a biological variable in our study.

### Structural modeling

The PyMOL Molecular Graphics System (Version 3.0 Schrödinger, LLC) was used to visualize the solved structure of mouse Rbpj bound to DNA (Protein Data Bank assembly 3BRG) (34,60). We used the PyMOL mutagenesis wizard to visualize and predict the impact of AOS-associated mutations, selecting the rotamer for each variant that occurs most frequently in proteins. Discs represent pairwise overlap of atomic van der Waals radii. The color and size of each disc correlates with the amount of overlap. All human residue numbers correspond to the numbering used in isoform Q06330-1.

### Protein purification

The pGEX-6P-1 plasmid construct encoding the highly conserved Rbpj core mouse residues 53-474 was used to generate each AOS variant through QuikChange site-directed mutagenesis using the primers listed in Table S3. DNA constructs were confirmed by Sanger sequencing, and proteins were purified as previously described (32,61). Protein concentrations were determined by measuring absorbance at 280 nm using a NanoDrop spectrophotometer. Protein purity was confirmed by SDS-PAGE with GelCode Blue staining (see Supplemental Figure S1B) per manufacturer’s protocol (Thermo Scientific Cat. #24590).

### Isothermal titration calorimetry (ITC)

ITC experiments were performed as previously described (32). Briefly, purified wild-type and variant Rbpj proteins were assessed for binding to the following ligands: A) the oligonucleotide sequence 5’–GGCAC<u>CGTGGGAA</u>ACTAGTG–3’, which contains a high-affinity RBPJ binding site (underlined); B) a peptide consisting of residues 1754-1781 of human NOTCH1, which includes the region of the NOTCH intracellular domain (NICD) that interacts with RBPJ; or C) human SHARP residues 2776-2833, which includes the region that binds RBPJ. The NOTCH1 peptide was synthesized as previously described (32), and human SHARP residues 2776-2833 were cloned into pSMT3 to produce recombinant protein with an N-terminal SMT3 and His tag as previously described (62). All proteins and DNA were dialyzed overnight in a buffer containing 50 mM sodium phosphate (pH 6.5) and 150 mM sodium chloride buffer. Experiments were done in triplicate using a MicroCal VP-ITC. The RBPJ + DNA experiments were conducted at 10°C, while the RBPJ + NICD/SHARP experiments were conducted at 25°C. All experiments were performed using 20 injections of 14 µL each. Heat of dilution experiments were conducted by injecting each ligand (DNA, NICD, or SHARP) in the syringe into a buffer-only solution in the cell. The heat of dilution experiment was subtracted from the experimental data before fitting. The raw data was analyzed using ORIGIN software and fit to a one-site binding model. A two-tailed t-test was used to compare wild-type Rbpj to each variant, with a p-value < 0.05 indicating a significant difference.

### Electrophoretic Mobility Shift Assays (EMSAs)

EMSAs were performed as described previously (32,63–65). In brief, the 5’– CGAACGAGGCAAACCTAGGCTAGAGGCACCGTGGGAAACTAGTGCGGGCGTGGCT–3’ oligonucleotide containing a single high-affinity Rbpj binding site (underlined) was annealed to a 5’IRDye-700-labeled oligonucleotide 5’–AGCCACGCCCGCACT– 3’ that is complementary to the 3’ end. This duplex DNA was filled in using DNA polymerase I and the double-stranded DNA was used as bait for the purified Rbpj wild-type and variant proteins. Binding reactions were incubated for 20 minutes at room temperature, and protein-DNA complexes were separated by acrylamide gel electrophoresis. Gels were run for 2 hours at 150V and then imaged using a LI-COR Odyssey CLx scanner. Band intensity was quantified using Image Studio^TM^ software (LI-COR Biotech LLC). Each experiment was performed in triplicate. A one-way ANOVA with Tukey post-hoc correction was used to compare wild-type Rbpj to each variant, with a p-value < 0.05 indicating a significant difference.

### Mice

Mice carrying *Rbpj^S358R^*, *Rbpj^E89G^*, and *Rbpj^E89G,flox^* alleles were generated in collaboration with the CCHMC Transgenic Animal and Genome Editing Facility (TAGE, RRID:SCR_022642) using CRISPR/Cas9 genome editing of CD1 mice unless otherwise noted. For the *Rbpj^S358R^* allele, we targeted cleavage to the appropriate site surrounding the S358 codon with the single guide RNA (sgRNA) 5’– TCCCTCATAGAACGTGTACTCGG–3’ and introduced the donor oligonucleotide 5’– ATCATTAGAACTGATAAAGCTGAGTATACG–3’, which both substituted an arginine in place of S358 and introduced a DdeI restriction site to facilitate genotyping. For the *Rbpj^E89G^* and *Rbpj^E89G,flox^* alleles, we targeted cleavage to the appropriate site surrounding the E89 codon with the sgRNA 5’–AGTCTTACGGAAATGAAAAACGG–3’ and introduced the donor oligonucleotide 5’–CAGAAGTCATATGGGAATGGAAAA–3’, which both substituted a glycine in place of E89 and introduced a NdeI restriction site to facilitate genotyping. The *Rbpj^E89G^* allele was made by editing wild-type CD1 mice, while the *Rbpj^E89G,flox^* allele was made in outbred mice with existing flox sites surrounding exons 6 and 7 of the *Rbpj* gene (35). The genotypes of founder animals were confirmed using Sanger sequencing by the CCHMC Genomics Sequencing Facility (RRID:SCR_022630).

The other mouse lines used in this study include three *N1* alleles: *N1^tm1Con^* (36) deletes genomic regions encoding amino acids 1056-2049, which includes several EGF repeats, the entire transmembrane region, and the Ankyrin repeats, and therefore is considered a constitutive null allele. We refer to this allele as *N1^null^*. The *N1^tm2Agt^* allele (37) was generated by incorporating loxP sites flanking the promoter and part of exon 1 followed Cre-mediated recombination in the germline to make a constitutive *N1* null allele that we refer to here as *N1^gKO^*. The *N1^tm2Rko^* allele (41) was independently made in the Kopan lab by inserting loxP sites in nearly identical sequences as Radtke et al., and when these loxP sites are recombined, they remove the promoter and exon 1. We refer to this conditional allele as *N1^flox^*. The other alleles used in this study were *Rbpj^null^* (66), *Rbpj^flox^* (35), *N2^LacZ^* (67), and *Tie2-Cre^Ywa^* (68). Offspring were genotyped using the methods and primers outlined in Table S2.

### Timed embryonic harvest

Gestation was timed such that observation of a vaginal plug was considered embryonic day 0.5 (E0.5). Pregnant dams were euthanized via CO_2_ inhalation followed by cervical dislocation, and the uterus was removed and placed into PBS on ice. Embryos were harvested from the uterus and imaged with a Nikon SMZ 1500 stereoscope prior to collection of required tissues. Specifically, the forelimbs, head, and/or heart were collected for analysis and placed into 4% paraformaldehyde (PFA) in PBS and incubated at 4°C overnight.

### Western blotting

Single E10.5 *Rbpj^+/+^* and *Rbpj^E89G,flox/E89G,flox^* embryos were homogenized in 2X Laemmli sample buffer for Western blot analysis. Samples were sonicated and stored at -80°C. Protein extracts (whole embryos for homozygotes, half embryos for wild-type controls) were run on a BIO-RAD 4-20% Mini-PROTEAN TGX Stain-Free Precast Gel (Cat. #456-8093) and transferred to a PVDF membrane via semi-dry transfer. The membrane was washed with water and then PBS before blocking with 0.5% casein in PBS for one hour at room temperature. The membrane was subsequently washed in PBS with 0.1% Tween-20, blocked in 0.5% Casein with 0.05% Tween-20 in PBS (pH 7.4) for one hour at room temperature, and then incubated in primary antibodies against Rbpj (1:1000, Cell Signaling Technology Cat. #5313) and beta-actin (1:2000, LI-COR Cat. #926-42212) overnight at 4°C. The membrane was then washed in PBS with 0.1% Tween-20 and incubated with secondary antibodies (1:4000 goat anti-rabbit IgG AF555, Invitrogen Cat. #A-21429 and 1:4000 donkey anti-mouse IgG 680RD, LI-COR Cat. #926-68072) at room temperature for 90 minutes. Finally, the membrane was washed in PBS with 0.1% Tween-20 and then imaged using a LI-COR Odyssey CLx scanner. Band intensity was quantified using the Image Lab Software Suite (BIO-RAD), and Rbpj was normalized to beta-actin levels.

### Embryonic and postnatal heart histochemistry

After overnight fixation in 4% PFA, E16.5 or postnatal hearts were washed 3 x 5 minutes in PBS and submitted to the Integrated Pathology Research Facility for processing and embedding in paraffin (RRID:SCR_022637). Hearts were then serially sectioned and stained with hematoxylin and eosin as described previously (69). Stained heart sections were imaged using a Nikon NiE upright widefield microscope.

### Embryonic yolk sac vascular assays

Embryos at 14.5 or 16.5 days of gestation were harvested from the uterus and imaged from multiple angles with a Nikon SMZ 1500 stereoscope prior to removal of the yolk sac. Yolk sac vasculature was considered ‘reduced’ if vitelline vessels were absent or markedly narrowed and/or if the capillary plexus extended over less than half of the yolk sac surface area. Yolk sac vasculature was scored by researchers blinded to genotype.

### Embryonic skin vascular assays

Embryonic skin vascular assays were performed essentially as previously described (70). In brief, PFA was gradually removed from E14.5 forelimbs and heads by washing 3 x 5 minutes in PBS. Tissues were transferred to 100% methanol (MeOH) for storage at -20°C. Using fine forceps, the skin was carefully removed from the forelimbs and heads and then rehydrated through a graded series of MeOH/PBT (PBS + 0.2% Triton X-100) washes. Skins were blocked with 10% normal donkey serum in PBT for 2 hours at room temperature and incubated with a primary antibody specific to endothelial cells (1:300 rat anti-CD31, BD Cat. #553369) overnight at 4°C. These skin preparations were then washed 5 x 15 minutes at room temperature with 2% normal donkey serum in PBT and incubated with a secondary antibody (1:300 donkey anti-rat AF647, Jackson ImmunoResearch Laboratories Inc. Cat. #712-605-153) for one hour at room temperature. Skins were again washed 5 x 15 minutes at room temperature. Skin samples were mounted on slides and imaged using a Nikon A1R inverted confocal microscope. Image analysis and quantification was performed with AngioTool (71) and Imaris (72) software.

### Statistics

Mouse viability was analyzed using the Chi squared (ꭓ^2^) test for deviation from expected Mendelian ratios. Fisher’s exact test was used to determine whether the frequency of a phenotype differed between groups. Additional statistical tests are as described in corresponding figure legends. For all statistic tests, [*] p < 0.05. [**] p < 0.01. [***] p < 0.001. [****] p < 0.0001, and NS = not significant.

### Study Approval

All animal experiments were carried out under protocols approved by the Institutional Animal Care and Use Committee (IACUC, #2016-0105 and #2021-0086) at Cincinnati Children’s Hospital Medical Center (CCHMC).

## Data Availability

All values underlying graphed data are available in the Supporting Data Values file.

## Author Contributions

Conceptualization: R.A.K., R.K., and B.G.

Formal analysis: A.F.S., K.P., B.C., and E.K.G.

Funding acquisition: B.G.

Investigation: A.F.S., K.P., B.C., R.H., P.G., Z.Y., B.B., G.M., H.N., E.K.G.

Methodology: A.F.S., K.P., B.C., Z.Y., and E.K.G.

Project administration: R.A.K., R.K., and B.G.

Resources: Z.Y., L.S., and R.A.K.

Supervision: E.K.G., R.A.K., R.K., A.F.S., and B.G.

Writing – original draft preparation: A.F.S. and B.G.

Writing – review and editing: A.F.S., K.P., B.C., E.K.G., L.S., R.K., and B.G.

## Acknowledgements

This work was funded by a Research Innovation Project grant from Cincinnati Children’s Hospital Medical Center and National Science Foundation grant #2114950 to B.G. We thank Mei Xin for providing *Tie2-Cre^Ywa^* mice, Katherine Yutzey for her expertise on heart morphology, and Elisa Boscolo and Yoh-suke Mukouyama for advice on embryonic vascular assays. This publication was made possible, in part, using the following Cincinnati Children’s Hospital Medical Center (CCHMC) shared core facilities: the Transgenic Animal and Genome Editing Facility [RRID:SCR_022642], the Integrated Pathology Research Facility [RRID:SCR_022637], the Genomics Sequencing Facility [RRID:SCR_022630], and the Bio-Imaging and Analysis Facility [RRID: SCR_022628]. Parts of the figures were created with BioRender.com as indicated.

