## Supplemental Figures and Tables for "A mouse model of Adams-Oliver Syndrome reveals defective Notch1 signaling in endothelial cells as a driver of pathogenesis"

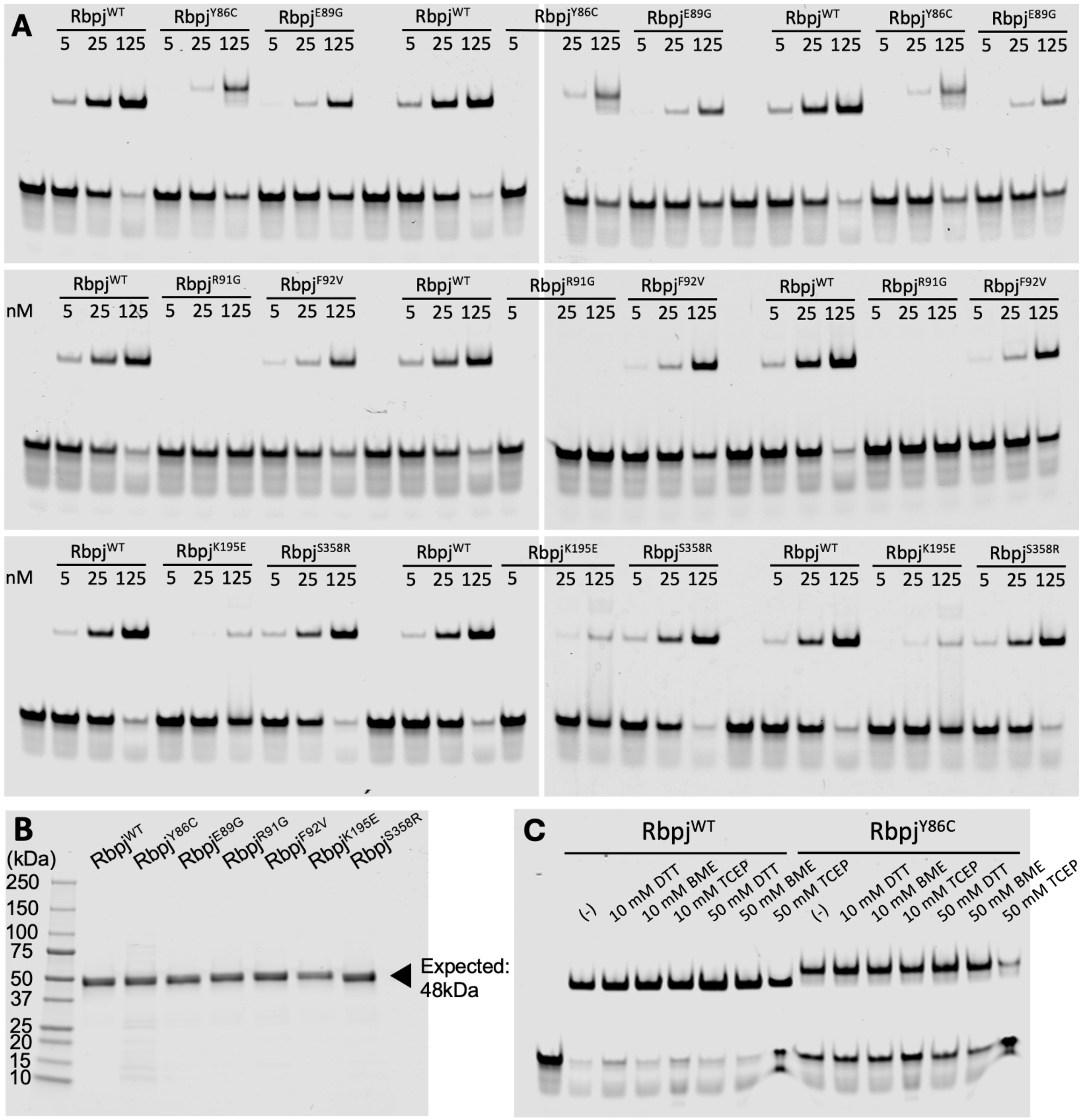


**Figure S1: DNA binding analysis of AOS-associated Rbpj variants using EMSAs.** A) Triplicate EMSAs of wild-type Rbpj and Rbpj variants Y86C, E89G, R91G, F92V, K195E, and S358R. The DNA probe contained a single high affinity Rbpj binding site (CGTGGGAA). Quantitative statistical analysis of probe depletion is presented in Figure 1I. B) SDS-PAGE with GelCode Blue staining of wild-type and variant Rbpj protein preparations shows a single 48 kDa band in each lane. C) EMSA of wild-type Rbpj and Rbpj variant Y86C (125 nM) with and without the indicated concentrations of reducing agents dithiothreitol (DTT), beta-mercaptoethanol (BME), and tris(2-carboxyethyl)phosphine (TCEP).

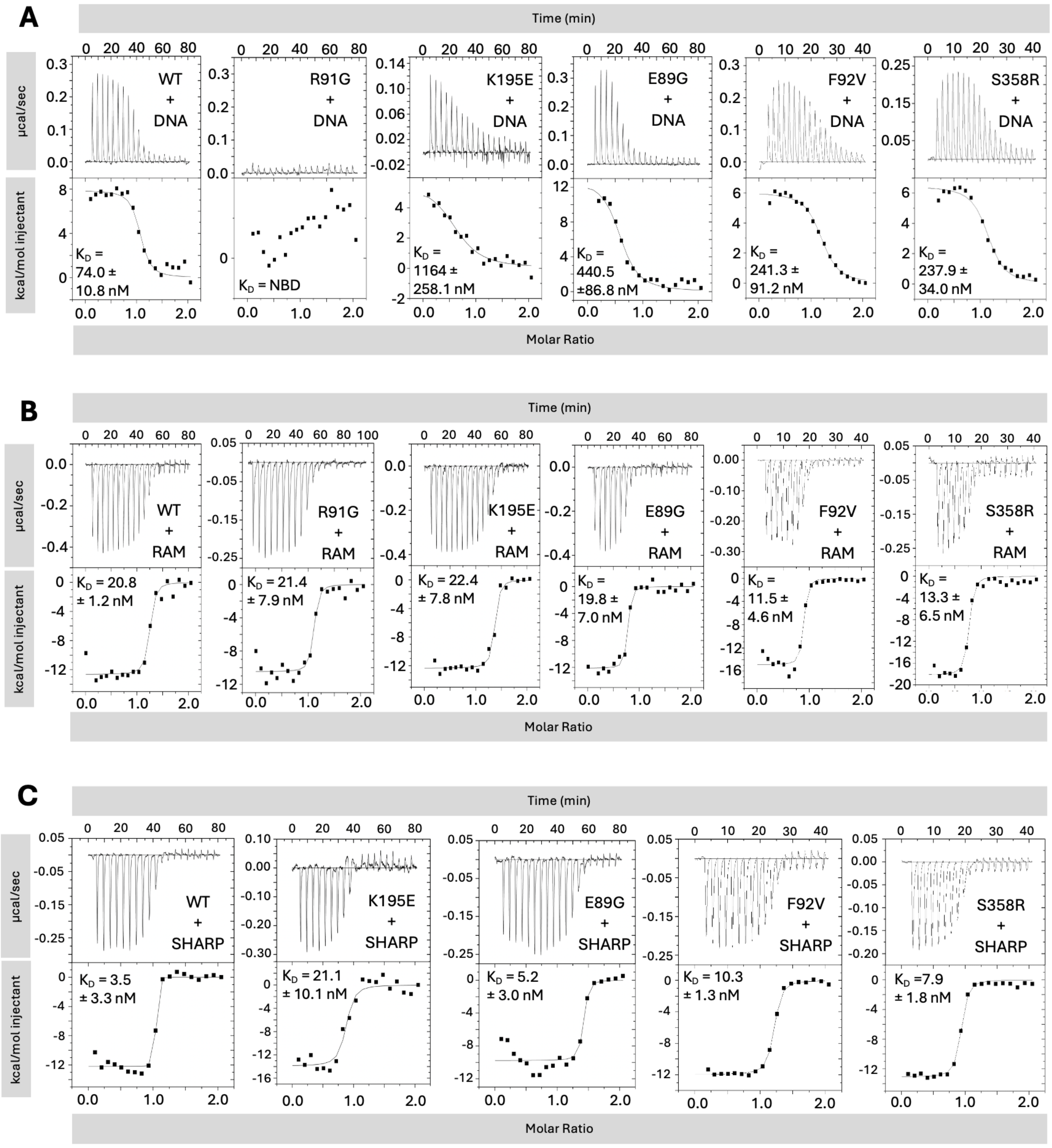


**Figure S2: Isothermal titration calorimetry studies show that AOS-associated Rbpj variants impair binding to DNA but not to co-factors.** Representative ITC thermograms showing the raw heat signal and nonlinear least squares fit to the data for mouse Rbpj variants binding to: A) an oligonucleotide containing a high-affinity Rbpj-binding site; B) the RBPJ association module (RAM) domain (VLLSRKRRRQHGQLWFPEGFKVSEASKK) of the human Notch1 intracellular domain; and C) a fusion protein of SMT3-SHARP 2776-2833 corresponding to the region of the human SHARP co-repressor protein that binds RBPJ. For experiments with wild-type (WT) Rbpj and the R91G, K195E, and E89G variants, the ligands (DNA, RAM, or SHARP) were placed in the syringe at 100μM and Rbpj protein was placed in the cell at 10μM. For experiments with the F92V variant, the ligand was placed in the syringe at 50μM and the Rbpj protein was placed in the cell at 5μM. For experiments with the S358R variant, the ligand was placed in the syringe at 40μM and the Rbpj protein was placed in the cell at 4μM. The mean dissociation constants ($K_D$) and standard deviations from triplicate experiments are reported. NBD = no binding detected. Statistical analysis of the DNA binding data is presented in Figure 1J and the statistical analysis of cofactor binding is presented in Table S1.

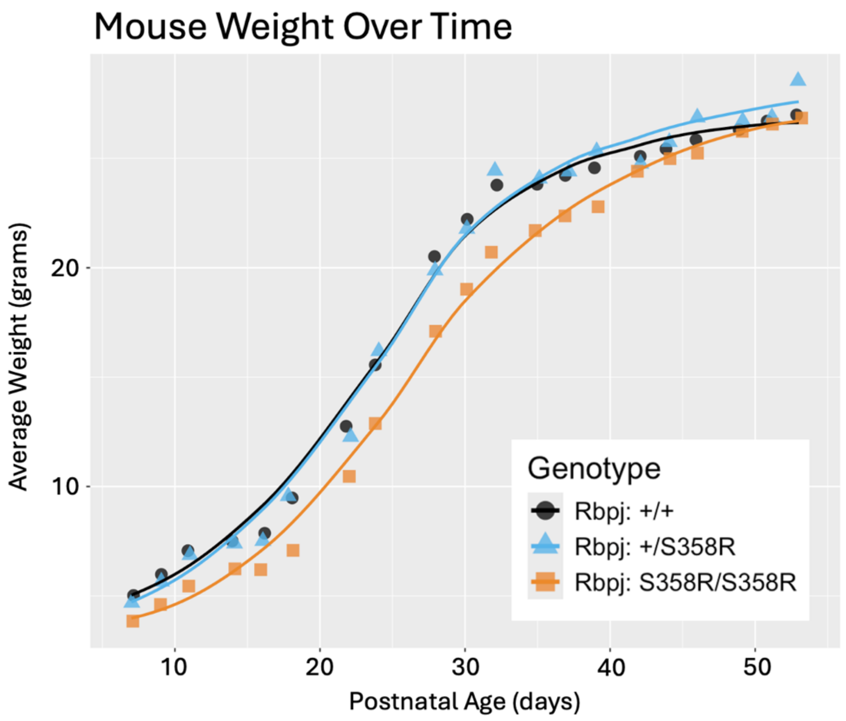


**Figure S3: *Rbpj*$^{S358R/S358R}$ mice exhibit early postnatal growth delay.** Plot of average body weight over time for *Rbpj*$^{+/+}$ (n = 13), *Rbpj*$^{+/S358R}$ (n = 25), and *Rbpj*$^{S358R/S358R}$ (n = 10) mice showing significant growth delay in *Rbpj*$^{S358R/S358R}$, but not *Rbpj*$^{+/S358R}$. Statistical analysis was performed using a two-way mixed ANOVA, which revealed that the interaction between genotype and postnatal age had a significant impact on weight (p = 2.09e-05). To further explore this interaction, we performed one-way ANOVAs at each timepoint with a Bonferroni correction for multiple comparisons and found weights were significantly different across genotypes until postnatal day 28. Pairwise comparisons from a Tukey post-hoc revealed that *Rbpj*$^{+/+}$ and *Rbpj*$^{S358R/S358R}$ were significantly different at these timepoints.

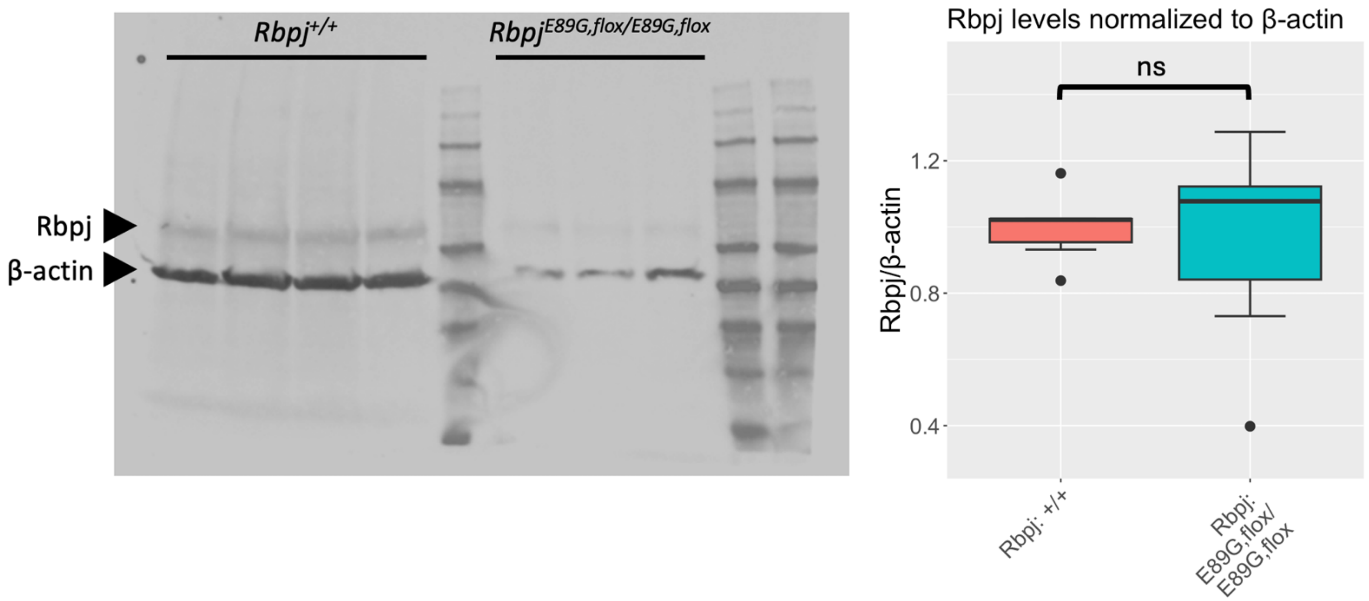


**Figure S4: Rbpj$^{E89G}$ is expressed at wild-type Rbpj levels.** Representative western blot of protein extracts from *Rbpj$^{+/+}$* (n = 6) and *Rbpj$^{E89G,flox/E89G,flox}$* (n = 8) embryos at E10.5 stained with antibodies against Rbpj and beta-actin (*left*). Box-and-whisker plot with quantification of Rbpj levels normalized to beta-actin (*right*). The limits of the box denote the interquartile range. A two-tailed t-test revealed no significant difference in Rbpj levels between genotypes.

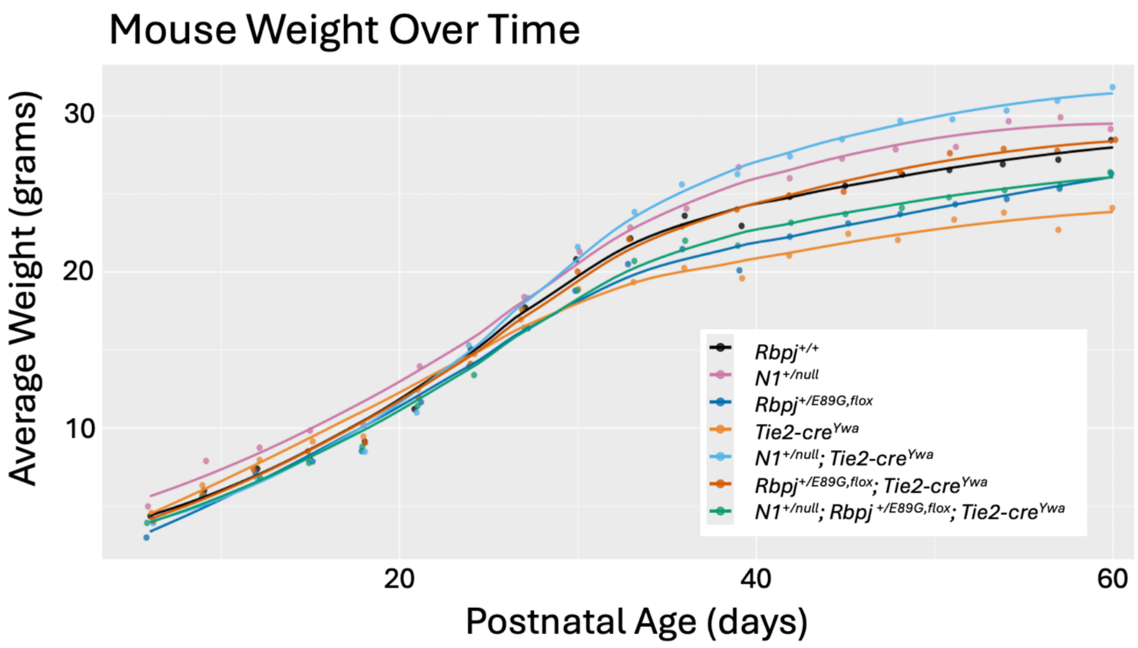


**Figure S5: *N1$^{+/null}$;Rbpj$^{+E89G,flox}$;Tie2-cre$^{Ywa}$* mice do not show postnatal growth delay.** Wild-type (n = 7), *N1$^{+/null}$* (n = 3), *Rbpj$^{+/E89G,flox}$* (n = 5), *Tie2-cre$^{Ywa}$* (n = 3), *N1$^{+/null}$;Tie2-cre$^{Ywa}$* (n = 5), *Rbpj$^{+/E89G,flox}$;Tie2-cre$^{Ywa}$* (n = 7), and *N1$^{+/null}$;Rbpj$^{+/E89G,flox}$;Tie2-cre$^{Ywa}$* (n = 9) mice were weighed every 3 days from postnatal day 6 (P6) through postnatal day 60 (P60). *N1$^{+/null}$;Rbpj$^{+/E89G,flox}$* mice could not be assessed due to insufficient numbers surviving postnatally. Two-way mixed ANOVA revealed that neither genotype nor the interaction between genotype and postnatal age had a statistically significant effect on weight.

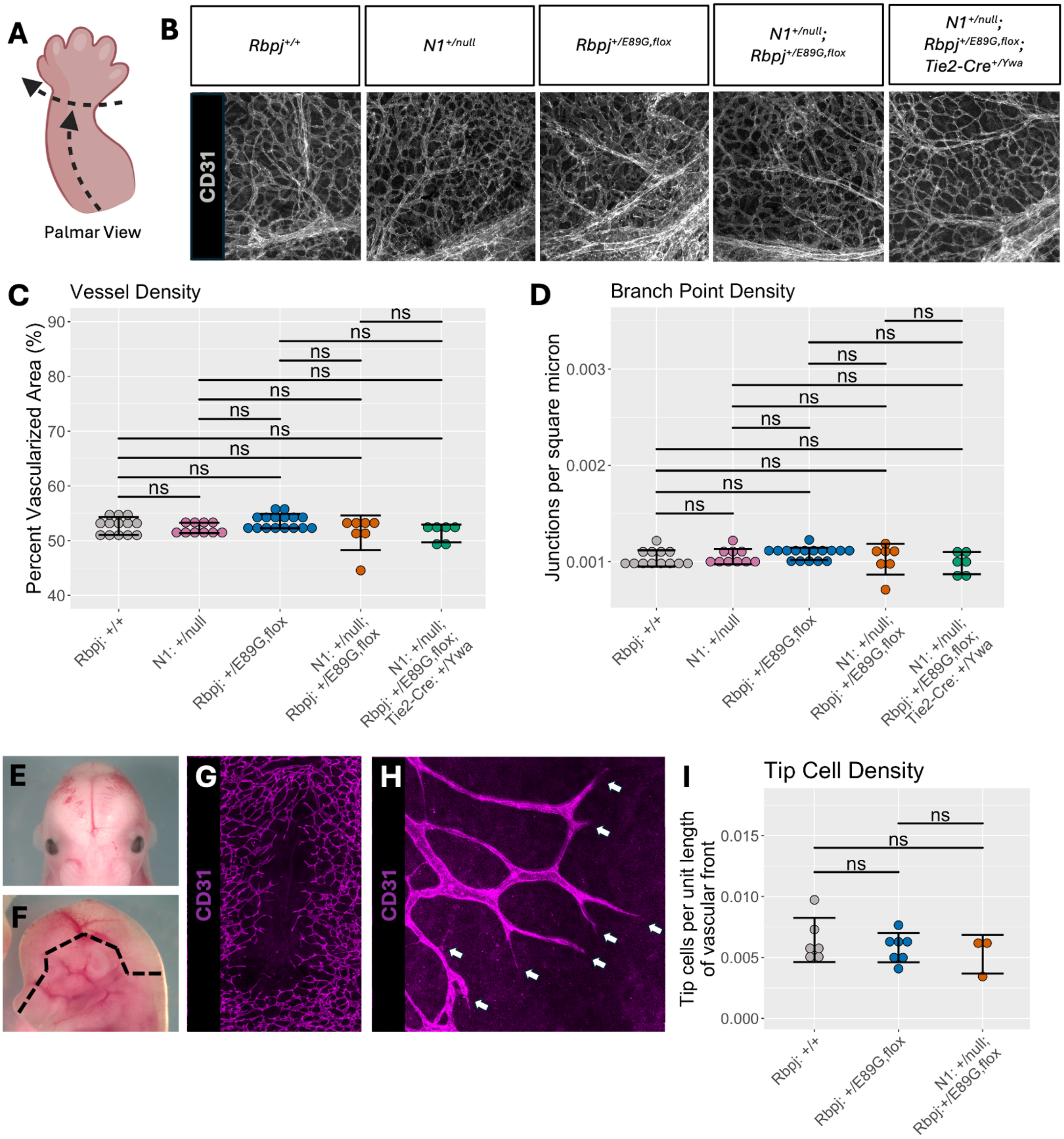


**Figure S6: Skin vascular assays show no difference between *N1$^{+/null}$; Rbpj$^{+/E89G,flox}$* embryos and littermates**. A) Schematic for removal of skin from forelimbs of E14.5 embryos by cutting from proximal to distal down the ventral side of the forelimb and circumferentially around the wrist. Created with *BioRender*. B) Representative images of E14.5 forelimb skin vasculature stained by CD31 at 20X magnification. C-D) Graphs showing the quantification of vessel density (C) and branch point density (D) in E14.5 forelimb skin vasculature. Groups were compared using two-way ANOVA followed by two-tailed t-tests with Bonferroni correction. E) Representative image of an E14.5 embryo with active vascular extension towards the midline scalp. F) Schematic for removal of scalps from E14.5 embryos by cutting along dashed lines on both sides of the head. G) Representative image of an E14.5 scalp with vessels stained with CD31 at 20X magnification. Note midline unvascularized area and active vascular extension towards the midline. H) Tip cells (white arrows) at the vascular front are identifiable by long, thin extensions. I) Graph showing the quantification of tip cell density at the scalp midline vascular front in E14.5 embryos. Two-tailed t-tests with Bonferroni correction showed no significant difference in tip cell density between genotypes. NS = not significant.

| Rbpj | Ligand | $K_D$ (nM) | Fold Change | P-value | Study |
|---|---|---|---|---|---|
| $Rbpj^{WT}$ | NICD | 20.8 ± 1.2 | N/A | N/A | Gagliani et al., 2022 |
| $Rbpj^{S358R}$ | | 13.3 ± 6.5 | ~0.6 | NS | This Study |
| $Rbpj^{F92V}$ | | 11.5 ± 4.6 | ~0.6 | NS | This Study |
| $Rbpj^{E89G}$ | | 19.8 ± 7.0 | ~1 | NS | Gagliani et al., 2022 |
| $Rbpj^{K195E}$ | | 22.4 ± 7.8 | ~1.1 | NS | Gagliani et al., 2022 |
| $Rbpj^{R91G}$ | | 21.4 ± 7.9 | ~1 | NS | This Study |
| $Rbpj^{WT}$ | Sharp | 3.5 ± 3.3 | N/A | N/A | Gagliani et al., 2022 |
| $Rbpj^{S358R}$ | | 7.9 ± 1.8 | ~2.3 | NS | This Study |
| $Rbpj^{F92V}$ | | 10.3 ± 1.3 | ~2.9 | NS | This Study |
| $Rbpj^{E89G}$ | | 5.2 ± 3.0 | ~1.5 | NS | Gagliani et al., 2022 |
| $Rbpj^{K195E}$ | | 21.1 ± 10.1 | ~6 | NS | Gagliani et al., 2022 |

**Table S1: Isothermal titration calorimetry studies show that AOS-associated Rbpj variants do not significantly impact binding affinity to cofactors.** Tabulated ITC binding data of Rbpj variants to the Rbpj association module of human Notch1 intracellular domain (NICD) and the corepressor Sharp. Fold change is calculated relative to wild-type Rbpj. A two-tailed t-test was used to compare the dissociation constant ($K_D$) of wild-type Rbpj to each variant. N/A = not applicable. NS = not significant.

| Gene Variant | | Primer Sequences | Annealing Temp. (°C) | Restriction Enzyme Digest | Product Size (bp) |
|---|---|---|---|---|---|
| **Notch1$^{gKO}$** | | | | | |
| | N1-F | CTGACTTAGTAGGGGGAAAAC | 64 | (none) | ~400 |
| | N1-R | AATCAGAGCGGCCCATTGTCG | | | |
| **Notch1$^{null}$** | | | | | |
| | N1-F | ACGATATCGTGGTGCATACGCTCCTGTGCG | 64 | (none) | ~400 |
| | N1-R | GTCAGTTTCATAGCCTGAAGAACG | | | |
| **Rbpj$^{E89G}$ and Rbpj$^{E89G,flox}$** | | | | | |
| | Rbpj-flox-F | TGTTTGCCACCAGAATCTGTTTGTTATTTGC | 68 | (none) | WT: 198<br>Floxed: 248 |
| | Rbpj-flox-R | AGGTACCTGGTACTAACTGTCTGGGACCG | | | |
| | Rbpj-E89G-F | AGCATTCCTCTCATCTGCTAGAG | 62 | NdeI | Uncut: 275<br>Cut (WT): 231/44<br>Cut (E89G): 164/67/44 |
| | Rbpj-E89G-R | TGAACCAAGGAATAGAACCAAGTC | | | |
| **Rbpj$^{S358R}$** | | | | | |
| | Rbpj-S358R-F | CTCAAGTAGGTGAATGTCACAGAG | 60 | DdeI | Uncut (WT): 286<br>Cut (S358R): 168/105/13 |
| | Rbpj-S358R-R | GGCAAGTTATAGCTCAGAACAGC | | | |
| **Rbpj$^{null}$** | | | | | |
| | Rbpj-null-F | TGGCACTGTTCAATCGCCTT | 60 | (none) | ~350 |
| | Rbpj-null-R | GAGGAAATTGCATCGCATTGTCTGAG | | | |
| **Tie2-Cre$^{Ywa}$** | | | | | |
| | Tie2-Cre-F | GGGAAGTCGCAAAGTTGTGAG | 59 | (none) | ~350 |
| | Tie2-Cre-R | TAACCAGTGAAACAGCATTGC | | | |

**Table S2: Genotyping information for mouse lines.** Primer sequences are listed 5' – 3'. F = forward, R = reverse, WT = wild-type, bp = base pairs.

| Purpose | Primer Sequences |
|---|---|
| Cloning | GGAGGATCCCCACCCAAACGACTCACTAG |
| | GGTCTCGAGTTATGGCTCTGGTGTGTAGGTG |
| Introduce Y86C | AAATCGTTTTTCATTTCCGCAAGACTTCTGTGCAACTTTTGCATG |
| | CATGCAAAAGTTGCACAGAAGTCTTGCGGAAATGAAAAACGATTT |
| Introduce E89G | GGAGGGCAAAAAAATCGTTTTCCATTTCCGTAAGACTTCTGTG |
| | CACAGAAGTCTTACGGAAATGGAAAACGATTTTTTTGCCCTCC |
| Introduce R91G | GGAGGAGGGCAAAAAAATCCTTTTTCATTTCCGTAAGAC |
| | GTCTTACGGAAATGAAAAAGGATTTTTTTGCCCTCCTCC |
| Introduce F92V | CAAGGAGGAGGGCAAAAAACTCGTTTTTCATTTCCGTAAG |
| | CTTACGGAAATGAAAAACGAGTTTTTTGCCCTCCTCCTTG |
| Introduce K195E | TCTTCAGTGACTGCTTCTTCTCGGAGGGTTTGGAGATGACC |
| | GGTCATCTCCAAACCCTCCGAGAAGAAGCAGTCACTGAAGA |
| Introduce S358R | GTATTCTGCCTTGTCTGTCCTAATGATTGTCCAGGAA |
| | TTCCTGGACAATCATTAGGACAGACAAGGCAGAATAC |

**Table S3: Primers used for site-directed mutagenesis.** Primer sequences are listed 5’ – 3’.
